# Hippocampal and amygdala subfield volumes in obsessive-compulsive disorder differ according to medication status

**DOI:** 10.1101/2023.03.28.534348

**Authors:** Ziphozihle Ntwatwa, Christine Lochner, Annerine Roos, Tatum Sevenoaks, Jack van Honk, Pino Alonso, Marcelo C. Batistuzzo, Sunah Choi, Marcelo Q. Hoexter, Minah Kim, Jun S. Kwon, David Mataix-Cols, José M. Menchón, Euripides C. Miguel, Takashi Nakamae, Mary L. Phillips, Carles Soriano-Mas, Dick J. Veltman, Nynke A. Groenewold, Odile A. van den Heuvel, Dan J. Stein, Jonathan Ipser

## Abstract

**Intro:** Although it has been suggested that the hippocampus and amygdala (HA) are involved in the neurobiology of obsessive-compulsive disorder (OCD), volumetric findings have been inconsistent. Furthermore, the HA consist of heterogenous anatomic units with specific functions and cytoarchitecture, and little work has been undertaken on the volumetry of these subfields in OCD.

**Methods:** T1-weighted images from 381 patients with OCD and 338 healthy controls (HCs) from the OCD Brain Imaging Consortium were segmented to produce twelve hippocampal subfields and nine amygdala subfields using FreeSurfer 6.0. We assessed between-group differences in subfield volume using a mixed-effects model adjusted for age, quadratic effects of age, sex, site, and whole HA volume. Given evidence of confounding effects of clinical characteristics on brain volumes in OCD, we also performed subgroup analyses to examine subfield volume in relation to comorbid anxiety and depression, medication status, and symptom severity.

**Results:** Patients with OCD and HCs did not significantly differ in HA subfield volume. However, medicated patients with OCD had significantly smaller hippocampal dentate gyrus (*p*_*FDR*_=0.042, *d*=-0.26) and molecular layer (*p*_*FDR*_=0.042, d=-0.29) and larger lateral (*p*_*FDR*_=0.049, *d*=0.23) and basal (*p*_*FDR*_=0.049, *d*=0.25) amygdala subfields than HCs. Unmedicated patients had significantly smaller hippocampal CA1 (p_FDR_=0.016, d=-0.28) than HCs. No association was detected between any subfield volume and OCD severity.

**Conclusion:** Differences in HA subfields between OCD and HCs are dependent on medication status, in line with previous work on other brain volumetric alterations in OCD. This emphasizes the importance of considering psychotropic medication in neuroimaging studies of OCD.

## 1. Introduction

Obsessive-compulsive disorder (OCD) is a common psychiatric disorder characterized by persistent intrusive thoughts (obsessions) or repetitive ritualistic overt or covert behaviours (compulsions), or both (APA, 2022). Typical obsessive thoughts concerning, contamination, harm, sexual or religious ideas, and exactness are accompanied by anxiety or distress, which may, in turn, incite compulsions such as excessive cleaning, checking, and, ordering and arranging, and counting (Goodman et al., 2014). OCD has a lifetime prevalence of 1.9% to 2.5% in the adult population with strong negative impact on occupational and social functioning (Ruscio et al., 2010). In many cases, OCD is comorbid with other disorders, including major depressive disorder (MDD) and anxiety disorders (Nazeer et al., 2020; Ruscio et al., 2010). Additionally, differences in symptom severity are likely to contribute to OCD heterogeneity (Mataix-Cols et al., 2005; Stein et al., 2019; van Oudheusden et al., 2020).

Neuroimaging studies suggest that OCD is associated with structural and/or functional changes in the cortico-striato-thalamo-cortical loops (CSTC) (Pauls et al., 2014; van den Heuvel et al., 2016). However, emerging evidence suggests that OCD involves additional brain circuits including the cerebellar, fronto-parietal, and fronto-limbic circuits (van den Heuvel et al., 2016). There has also been interest in investigating the hippocampal formation and amygdala in OCD, given the established roles of these brain structures in anxiety (Brühl et al., 2014; González-García & Visser, 2023; Shi et al., 2023) and fear conditioning (Cheng et al., 2003). Indeed, an fMRI study suggested that during fear conditioning, the hippocampus has reduced activation in patients with OCD compared to healthy controls (HCs) (Milad et al., 2013), and a meta-analysis indicated increased amygdala activation during emotional processing in patients with OCD versus HCs (Thorsen et al., 2018).

However, structural MRI studies in OCD have yielded inconsistent findings, reporting both increases and decreases in HA volumes (Kwon et al., 2004; Pujol et al., 2004; Atmaca et al., 2008; Rao et al., 2018). Such inconsistency could be attributable to small sample sizes, the presence of comorbidities, or medication use. Additionally, previous studies investigated whole HA volumes rather than investigating subfield volumes, which may not reveal subtle OCD-related differences in volume when these vary between the individual subfields. Work from the Enhancing Neuroimaging Genetics through Meta-Analysis (ENIGMA) OCD Consortium (ENIGMA-OCD), a large brain imaging consortium, found smaller hippocampal volumes in medicated patients with OCD compared to HCs, which was also related to adult-onset OCD and comorbid depression (Boedhoe et al., 2017). These findings were corroborated by work from the multisite OCD Brain Imaging Consortium (OBIC) which demonstrated that smaller hippocampi were associated with medication use (Fouche et al., 2017; Fouche et al., 2022).

The HA are anatomically complex structures, consisting of multiple interconnected nuclei that can be segmented according to their cytoarchitecture, histochemistry and connectivity profile (Sah et al., 2003), -however little is known about hippocampal and amygdala subfield volumes in OCD. Recent developments in structural MRI segmentation techniques have allowed for the robust delineation of HA subfields using a Bayesian algorithm that is based on the transformation of manual segmentation to an automated atlas (Saygin et al., 2017). Indeed, a previous study showed that paediatric patients with OCD have larger hippocampal subfields, i.e., the left subiculum body, left cornu ammonis (CA) 4, left granule cell layer of dentate gyrus, left molecular layer, and right parasubiculum, compared to HCs (Vattimo et al., 2021). Recent reports indicate that medication-free patients with OCD have smaller volumes in the hippocampal subiculum, presubiculum, CA2/3 and tail (Zhang et al., 2019) and smaller basolateral and central amygdala subfield volumes (Zhang et al., 2020). However, these studies were conducted in smaller sample sizes, and did not include patients with psychotropic medication use.

To the best of our knowledge, this is the first study to investigate HA subfield volume in a large international multi-site sample of adult patients with OCD (n=381) and HCs (n=338). We utilized the automated segmentation algorithm on T1-weighted MRI scans to segment the HA into 12 and 9 subfields, respectively. Given the evidence of the potential confounding effects of clinical characteristics on brain volumes, we performed separate analysis for patients with and without psychotropic medication use and studied the effect of comorbid anxiety and depression. We also studied the association of subfield volumes with OCD symptom severity.

## 2. Methods

### 2.1. Participants and MRI acquisition

Sociodemographic and neuroimaging data were obtained from six research sites as part OBIC. Collaborating sites and participant details have been described in-depth in a previous publication (De Wit et al., 2014). Briefly, patients with OCD were recruited through local outpatient clinics, whereas HCs were sourced through local advertisements. All participants were screened for DSM-IV Axis I disorder. For the patient group, the primary diagnosis had to be OCD, but comorbidity with mood and anxiety disorders was allowed. To be included, healthy participants were required to be without current Axis I psychiatric disorders. Participants were excluded if they were younger than 18 or older than 65 years, had a current psychotic disorder, history of substance use disorder, intellectual disability and severe organic or neurological pathology. Additional data was collected on age of OCD onset, medication status and symptom severity (assessed with the Yale-Brown Obsessive Compulsive Scale (Y-BOCS)) (Goodman et al., 1989) (see Table 1A). Approval was obtained per site from all local ethical review boards and written informed consent was provided by each participant. In addition, for multi-site pooling of data, approval was obtained from the medical ethical committee Amsterdam, University Medical Center (UMC).

**Table 1A:** Demographic and clinical characteristics of patients with OCD and healthy controls.

|  | OCD (n=381) |  | HCs (n=338) |  | Statistical analysis |  |  |
| --- | --- | --- | --- | --- | --- | --- | --- |
| Characteristics | mean | SD | mean | SD | t | df | p |
| Age (years) | 32 | 9.4 | 30.2 | 9.3 | -2.5 | 708.7 | 0.001 |
| Education level (years) | 13.7 | 2.8 | 14.6 | 3.4 | 3.6 | 653 | < 0.001 |
| Y-BOCS score mean<sup>a</sup> | 25 | 6.2 | -- | -- |  |  |  |
| Age at onset of OCD<sup>b</sup> | 20.3 | 8.8 | -- | -- |  |  |  |
| Total hippocampal volume | 3544.4 | 340.4 | 3587.5 | 361.6 | 1.6 | 694.4 | 0.101 |
| Total amygdala volume | 1770.3 | 187.9 | 1783 | 198.1 | 0.8 | 696.3 | 0.377 |
|  | **N** | **%** | **N** | **%** |  |  |  |
| Male | 186 | 48.8 | 179 | 52.9 | -1.1 | 707 | 0.268 |
| Right-handed | 327 | 85.8 | 303 | 89.6 |  |  |  |
| Medication use at time of scan | 161 | 43.6 | -- | -- |  |  |  |

<sup>a</sup> Measured by the Yale-Brown Obsessive Compulsive Scale (YBOCS) total score. <sup>b</sup> As measured by the Y-BOCS symptom checklist.

### 2.2. MRI image analysis and segmentation

All participants underwent 1.5-T structural T1-weighted MRI scanning (for scan parameters and scan sequences see De Wit et al., 2014). MRI image analysis was performed on the high-performance computing (HPC) cluster at the University of Cape Town, South Africa. First, we applied the standard FreeSurfer v5.3 analysis pipeline using *recon-all* to initiate all cortical reconstruction processes http://surfer.nmr.mgh.harvard.edu/). *Recon-all* initiates bias-field correction to the T1-weighted images, as well as registration to Talairach space, intensity normalisation and skull stripping (Fischl et al., 2002).

Next, subfield segmentation was performed using the *segmentHA_T1*.*sh* script in FreeSurfer v6.0. This script simultaneously segments the HA, thereby preventing structural overlap (Iglesias et al., 2015). The probability atlas applied by the script is based on the transformation of *ex vivo* manual segmentation to an automated algorithm that segments *in vivo* MRI data in target space. The atlas was built using Bayesian inference based on a tetrahedral mesh spanning the amygdala and neighbouring structures (Saygin et al., 2017). For each participant the model produces left and right volumes for the HA subfields as well as whole HA volume and intracranial volume (ICV).

The hippocampus was segmented as follows: parasubiculum, presubiculum, subiculum, cornu ammonis (CA) sectors, CA1, CA2-3, CA4, dentate gyrus (DG), molecular layer (ML), hippocampus–amygdala transition area (HATA), fimbria, hippocampal tail, and hippocampal fissure (Iglesias et al., 2015). The hippocampal subfields were grouped according to the FreeSurfer 6.0 hippocampal module without head/body subdivision and the ML was not absorbed to the nearest DG layer (see https://surfer.nmr.mgh.harvard.edu/fswiki/HippocampalSubfieldsAndNucleiOfAmygdala). The amygdala was segmented into 7 nuclei (lateral (LA), basal (BA), accessory basal (AB), central (Ce), medial (Me), cortical (Co), paralaminar nucleus (PL)) and 2 transition areas (anterior amygdaloid area (AAA) and cortico-amygdaloid transition (CAT)). Schmitz-Koep et al., 2021 suggests that the amygdala can be grouped in the following three regions: basolateral ((BLA) lateral, basal, accessory basal, paralaminar nucleus), centro-medial (central and medial), and superficial area ((SFA) cortical, AAA and CAT). Shown in Figure 1 is an example of HA Freesurfer segmentation.

**Figure 1:**
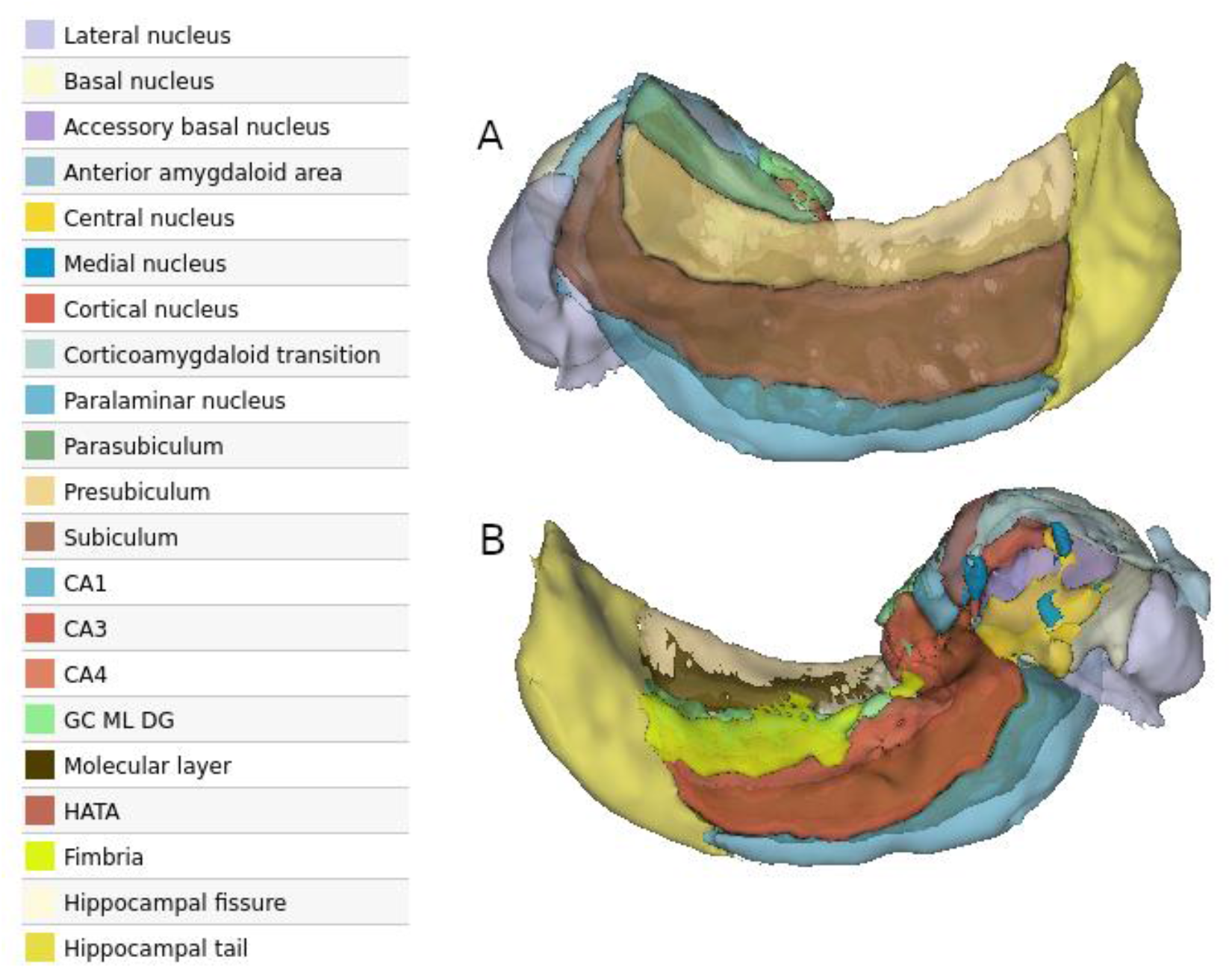
Visualisation of amygdala/hippocampal Freesurfer subfield segmentation from right hemisphere of single representative healthy control, using 3DSlicer (https://www.slicer.org/). A: Lateral view, B: Medial view. The hippocampal fimbria was not included in our analysis (Brown et al., 2020).

### 2.3. Quality control: visual inspection

We used a combination of visual inspection and quantitative measures to identify inaccurate subfield segmentation. To visually identify segmentation failures, we used an adaptation of the ENIGMA Consortium Quality Control protocol for subcortical and hippocampal subfields (https://enigma.ini.usc.edu/protocols/imaging-protocols/). In brief, three independent raters (ZN, AR, TS) examined each scan, slice-by-slice, within a html-based image gallery for partial or atypical segmentation. A list of questionable cases was generated for 3D inspection, using the Freeview utility included with FreeSurfer (Sämann et al., 2022). Additional cases were identified as follows: 1) we z-standardized each subfield and excluded participants whose score exceeded ± 5 SD from the mean (van der Meer et al., 2018, supplemental data): and 2) we generated automated outliers using a *R* script provided by ENIGMA-MDD working group (https://enigma.ini.usc.edu/ongoing/enigma-hippocampal-subfields). For the latter, participants flagged as outliers for 5+ subfields were added to the list for 3D inspection.

As shown in Table 2, 55 participants were excluded from the main analysis, i.e., 40 participants from visual QC, an additional 9 participants based on both visual QC after outlier flags, and 6 participants as their z-scores exceeded ±5SD from the mean of any subfield. The final sample consisted of 381 patients with OCD and 338 HCs.

**Table 2B:** Demographic and clinical characteristics of subgroup patients with OCD.

|  | **A: OCD on medication**<br>**(n=161)** | **B: OCD not on medication**<br>**(n=220)** | **C: OCD with anxiety**<br>**(n=25)** | **D: OCD without anxiety**<br>**(n=356)** | **E: OCD with MDD** | **F: OCD without MDD**<br>**(n=286)** | **G: HCs** | **Statistics** |
| --- | --- | --- | --- | --- | --- | --- | --- | --- |
|  | **Mean (SD)** | **Mean (SD)** | **Mean (SD)** | **Mean (SD)** | **Mean (SD)** | **Mean (SD)** |  |  |
| **Age (years)** | 32.89 (9.13) | 31.33 (9.54) | 34.48 (9.25) | 31.81 (9.39) | 34.99 (9.56) | 30.99 (9.14) | 30.2 (9.3) | A>G<br>C>G<br>D>G |
| **Education level (years)** | 12.90 (2.88) | 14.33 (2.62) | 14.83 (2.90) | 13.63 (2.81) | 13.48 (2.92) | 13.78 (2.79) | 14.6 (3.4) | A>G<br>D>G<br>E>G<br>F>G |
|  | **N (%)** | **N (%)** | **N (%)** | **N (%)** | **N (%)** | **N (%)** |  |  |
| **Male** | 83 (51.55) | 103 (46.81) | 7 (28) | 179 (50.28) | 32 (33.68) | 154 (53.84) | 179 (52.9) | C>G<br>E>G |

**Note:** mean age of OCD on medication (A), OCD with anxiety (C), OCD without anxiety (D) significantly larger than HCs (G), respectively. Years of education significantly greater in OCD on medication (A), OCD without anxiety (D), OCD with MDD (E), without MDD (F) significantly larger than HCs (G), respectively. Significantly larger number of men in OCD with anxiety (C), OCD with MDD (E) than HCs (G), respectively.

### 2.4. Statistical analysis

#### 2.4.1. Covariate selection

In addition to adjustment for age, quadratic effects of age, sex, and scanner site across all analyses (Barnes et al., 2010; Chen et al., 2018; Nordenskjöld et al., 2013; Sargolzaei et al., 2015), we also included those covariates that demonstrated an association with HA volumes in specific models (see supplemental data).

#### 2.4.2. Linear mixed-effects models

All statistical analyses were conducted in R (https://www.r-project.org/). We used the R package *lme4* to perform our analysis and outputted mixed-effects *d* effect sizes, as calculated using the t values from linear mixed effects models (equation 22, Nakagawa & Cuthill, 2007) that included a random intercept for scan site. We averaged the left and right hemisphere volumes to produce a single value per participant (Boedhoe et al., 2017). In total, tests were performed for 21 separate subfields. All analysis were corrected for multiple comparison across 21 subfields using the false discovery rate (FDR). We corrected all models for the total subfield volume using the combined HA volume (as recommended in the FreeSurfer manual; https://surfer.nmr.mgh.harvard.edu/fswiki/HippocampalSubfieldsAndNucleiOfAmygdala). We performed separate analysis where we compared subgroups of interest to HCs, including patients with OCD without anxiety comorbidity (n=356), those with of MDD (n=95), those without MDD (n=286), those with a history of psychotropic medication use (n=161), and those without medication use (n=220). We did not include patients with OCD with anxiety comorbidity (n=25) due to low sample size.

## 3. Results

### 3.1. Sample characteristics

In the full sample (OCD: n=381; HC: n=338, see Table 1A), patients with OCD were significantly older (OCD: 32.0 years (SD = 9.4); HCs: 30.2 years (SD =9.3), t = -2.5, p = 0.012) and on average, had lower education level (OCD: 13.7 years (SD=2.8); HCs (14.6 years (SD =3.4), t = 3.6, p < 0.001) than HCs. There were no significant differences in sex and whole HA volume between patients with OCD and HCs. The mean Y-BOCS score for patients with OCD was 24.9 (SD=6.2). See Table 1B for details on demographic and clinical characteristics of subgroup patients with OCD.

### 3.2. Group difference in subfield volumes

Between group comparisons were conducted on 381 patients with OCD and 338 HCs. There were no significant differences in HA subfield volumes (p < 0.05, FDR-corrected), after adjusting for age, quadratic effects of age, sex, site, and whole HA volume, between patients with OCD and HCs (Figure 2A/Table 3). There were no group by age interaction effects for all subfield volumes.

**Table 3:**
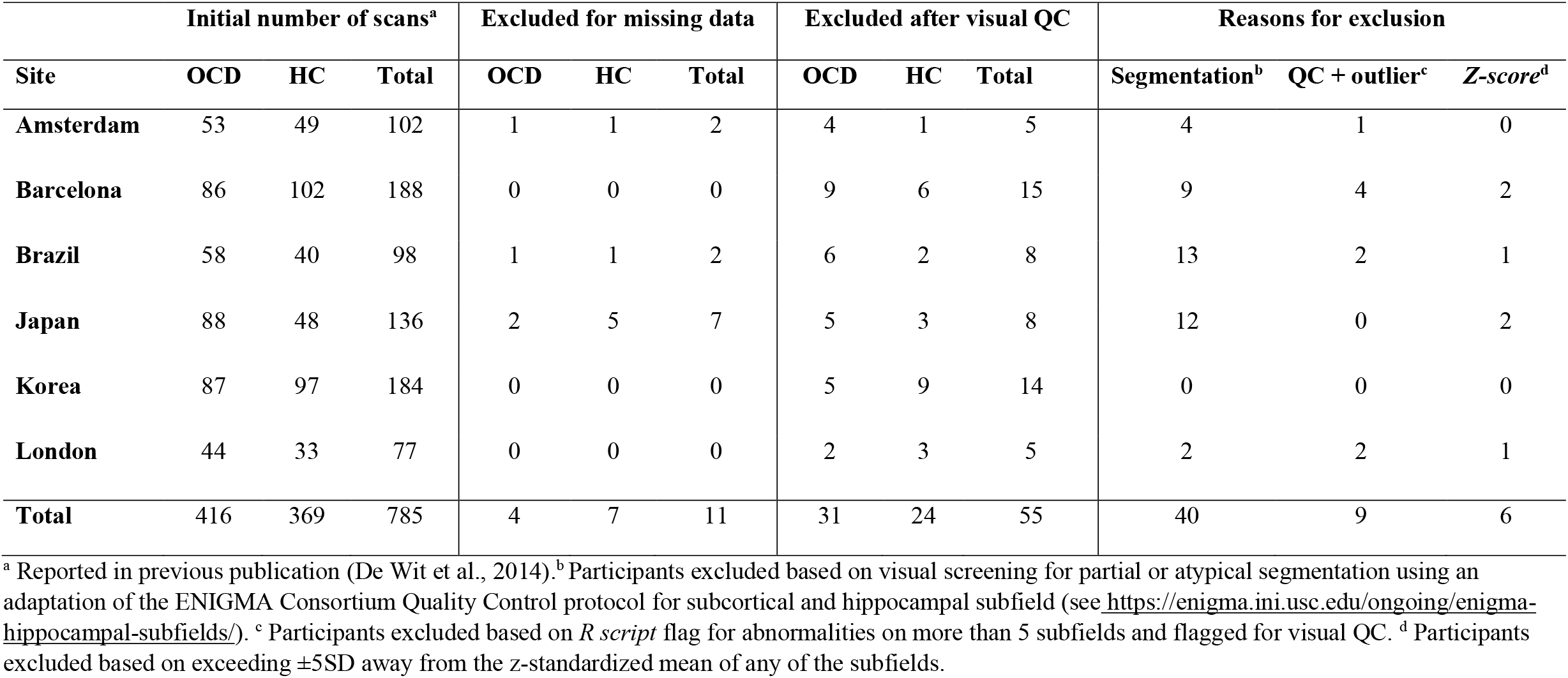
Number of scans provided and excluded for patients with OCD and healthy controls after quality checking.

**Table 4:** Mixed effect size estimates (d), SE, uncorrected and corrected (FDR) p-value for hippocampal and amygdala subfield volumes between patients with OCD (n=381) and healthy controls (n=338).

|  | Effect size (mixed-effects<br>*d*) | Standard Error<br>(SE) | P value<br>(uncorrected) | P value (corrected<br>(*p*<sub>FDR</sub>)) |
| --- | --- | --- | --- | --- |
| **Hippocampal subfields** |  |  |  |  |
| Parasubiculum | 0.051 | 0.075 | 0.443 | 0.664 |
| Presubiculum | 0.152 | 0.075 | 0.037 | 0.156 |
| Subiculum | 0.039 | 0.075 | 0.540 | 0.756 |
| CA1 | -0.196 | 0.075 | 0.006 | 0.063 |
| CA3 | -0.096 | 0.075 | 0.164 | 0.353 |
| CA4 | -0.111 | 0.075 | 0.104 | 0.353 |
| DG | -0.155 | 0.075 | 0.023 | 0.150 |
| ML | -0.207 | 0.075 | 0.003 | 0.060 |
| HATA | -0.049 | 0.075 | 0.363 | 0.636 |
| fimbria | 0.108 | 0.075 | 0.122 | 0.353 |
| Hippocampal fissure | 0.012 | 0.075 | 0.864 | 0.955 |
| Hippocampal tail | 0.030 | 0.075 | 0.681 | 0.841 |
| **Amygdala subfields** |  |  |  |  |
| Lateral nucleus | 0.095 | 0.075 | 0.168 | 0.353 |
| Basal nucleus | 0.102 | 0.075 | 0.149 | 0.353 |
| Accessory basal nucleus | 0.017 | 0.075 | 0.802 | 0.935 |
| Central nucleus | 0.036 | 0.075 | 0.631 | 0.828 |
| Medial nucleus | -0.076 | 0.075 | 0.296 | 0.564 |
| Cortical nucleus | 0.004 | 0.075 | 0.956 | 0.979 |
| Paralaminar nucleus | 0.154 | 0.075 | 0.029 | 0.150 |
| CAT | 0.002 | 0.075 | 0.979 | 0.979 |
| AAA | -0.050 | 0.075 | 0.419 | 0.664 |

Abbreviations: cornu ammonis (CA) sectors, CA1, CA2-3, CA4, granule cell layer of dentate gyrus (DG), molecular layer (ML), hippocampus–amygdala transition area (HATA), corticoamygdaloid transition area (CAT), anterior amygdaloid area (AAA). *P*-values are presented before and after correction for multiple comparison using the false discovery rate (FDR). Data presented with SE.

**Figure 2A:**
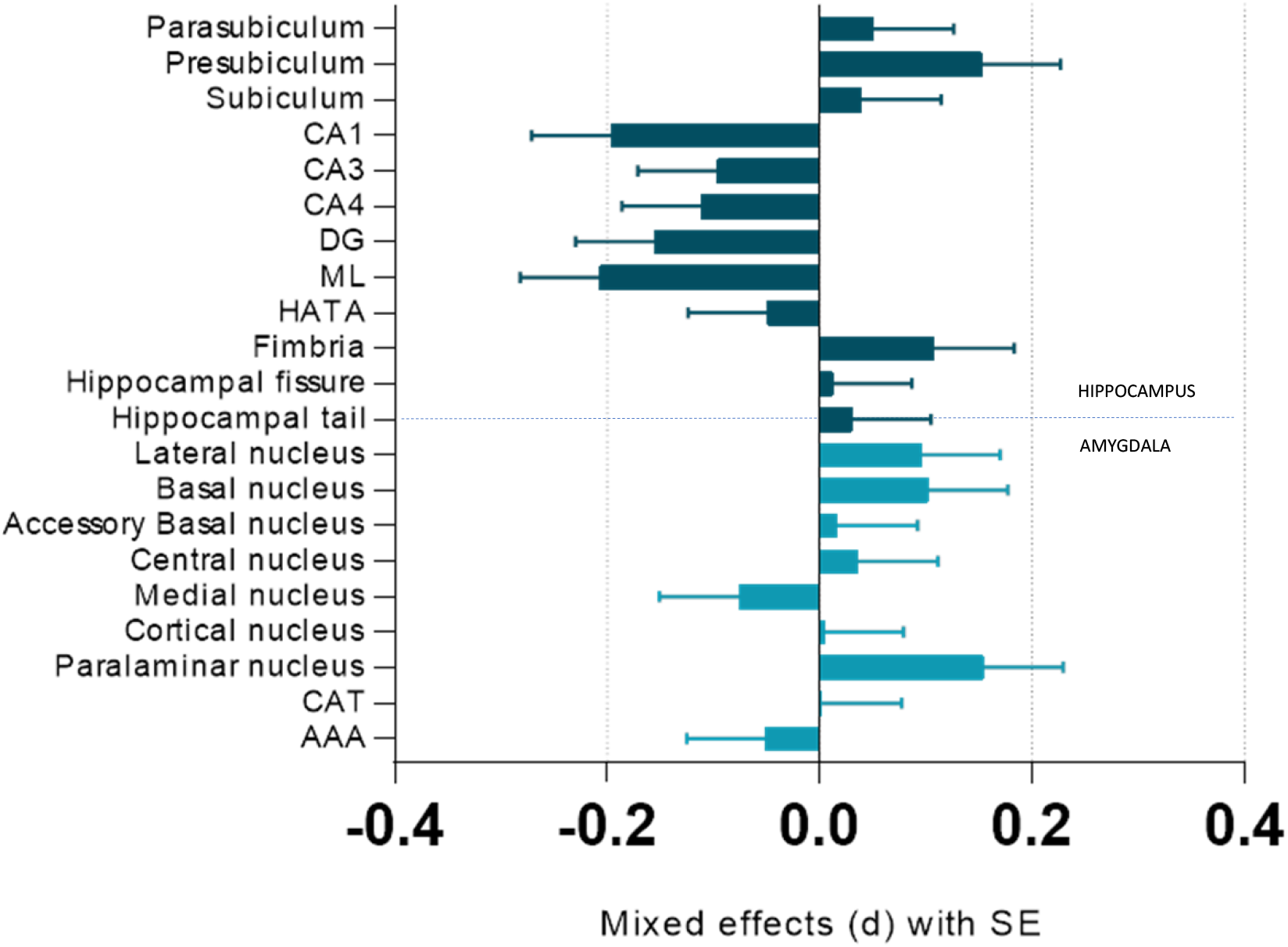
Mixed effect size estimates (d) for hippocampal and amygdala subfield volumes between patients with OCD (n=381) and healthy controls (n=338). Data presented with SE. Hippocampal subfields presented in dark blue, amygdala subfields presented in light blue. Abbreviations: cornu ammonis (CA) sectors, CA1, CA2-3, CA4, granule cell layer of dentate gyrus (DG), molecular layer (ML), hippocampus–amygdala transition area (HATA), corticoamygdaloid transition area (CAT), anterior amygdaloid area (AAA).

### 3.3. Subgroup analysis: association of subfields volume and clinical factors

#### 3.3.1. Psychotropic medication

Medicated patients with OCD (n=161) had significantly smaller hippocampal DG (*p*_*FDR*_= 0.042, *d*= -0.26) and ML (*p*_*FDR*_= 0.042, *d*= -0.29), and larger LA (*p*_*FDR*_= 0.049, *d*= 0.23) and BA (*p*_*FDR*_= 0.049, *d*= 0.25) amygdala, compared to HCs (n=291) (Figure 2B). Unmedicated patients with OCD (n=220) had significantly smaller hippocampal CA1 subfield volumes (*p*_*FDR*_= 0.016, d= -0.28) than HCs (n=338).

**Figure 2B:**
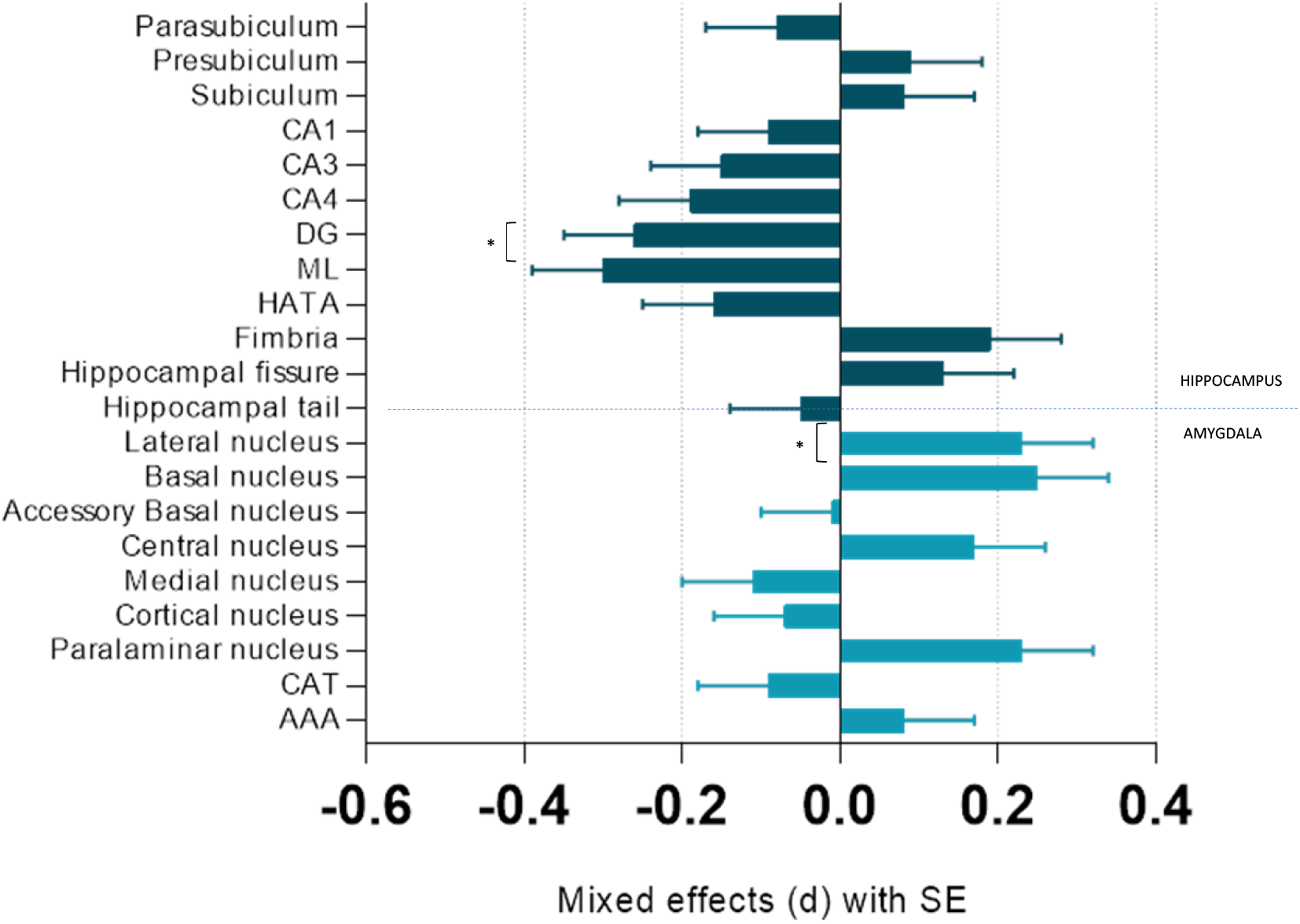
Mixed effect size estimates (d) for hippocampal and amygdala subfield volumes between **medicated patients with OCD** (n=161) and healthy controls (n=291). Data presented with SE, (*) Denotes significant difference *p*<0.05. Hippocampal subfields presented in dark blue, amygdala subfields presented in light blue. Abbreviations: cornu ammonis (CA) sectors, CA1, CA2-3, CA4, granule cell layer of dentate gyrus (DG), molecular layer (ML), hippocampus–amygdala transition area (HATA), corticoamygdaloid transition area (CAT), anterior amygdaloid area (AAA).

#### 3.3.2. Anxiety and MDD comorbidity

There were no significant differences in HA subfield volumes between HCs (n=338) and either patients with OCD with (n=95) or without (n=286) MDD, nor patients with OCD without anxiety comorbidity (n=356).

#### 3.3.3. Symptom severity

We also tested whether subfield volumes were influenced by OCD symptom severity. We found no significant association between the volume of subfields and Y-BOCS scores (see supplemental data).

## 4. Discussion

In this paper we report findings from the largest neuroimaging study of HA subfield volumes in OCD conducted to date. While we did not detect any significant differences between patients with OCD and HCs in HA subfield volumes after multiple comparison correction, one key finding emerged from our analysis. We found that, compared to HCs, medicated patients with OCD had both smaller volumes in the DG and ML of the hippocampal formation and larger volumes in the LA and BA amygdala. Our findings affirm previous work demonstrating medication effects on subcortical brain volumes in OCD, suggesting that medication status is a robust confounding factor that may influence the ability to detect neuroanatomical abnormalities in OCD (Boedhoe et al., 2017; van den Heuvel et al., 2022; Weeland et al., 2022).

Our finding, that the hippocampal subfield DG and ML were significantly smaller in medicated patients with OCD than in HCs, is consistent with previous ENIGMA-OCD studies showing smaller hippocampi in medicated patients with OCD (Boedhoe et al., 2017; Fouche et al., 2017; Ivanov et al., 2022). However, our findings contradict studies showing smaller volumes in the hippocampal subiculum, presubiculum and tail in patients with OCD compared to HCs, although these studies were performed in smaller sample sizes and did not include clinical characteristics (Zhang et al., 2019). In rodent studies, the DG supports hippocampal-based neurogenesis which in turn influences hippocampal plasticity (Malberg et al., 2000; Mandyam, 2013; Perera et al., 2007; Wang et al., 2008), and there is evidence that antidepressants increase proliferation in hippocampal-based neurogenesis (McEwen, 1999; Santarelli et al., 2003). These findings are in contrast to our observation of smaller DG and ML hippocampal volumes in patients with OCD.

Although the hippocampus has been commonly studied in relation to adult neurogenesis (Shapiro et al., 2009), some evidence shows that the human amygdala may be involved in postnatal neurogenesis with cell turnover rates that are comparable to the hippocampus (Roeder et al., 2022). Rodent work indicates that the LA and BA contain immunoreactive neural cell adhesion molecules that could allow for the amygdala to participate in neuronal plasticity (Nacher et al., 2002). Additional work in rodent and non-human primates demonstrate that antidepressant-modulated neurogenesis enhances neuronal and glial cell growth in the amygdala (Bernier et al., 2002; Fowler et al., 2002). Although these findings may explain the observation of the larger amygdala BA and LA subfields in our medicated patients with OCD, it is not known whether the subfield volumetric differences observed in medicated patients in the present study reflect an innate response to medication use or neurotoxic effect of medication. Further investigation is required to elucidate the effects of medication in subcortical volumes in OCD.

Another finding was that unmedicated patients with OCD had smaller hippocampal CA1 volume, compared to HCs. Although the CA1 is shown to be susceptible to stress-induced atrophy (Bartsch et al., 2015; Kassem et al., 2013), it is unclear whether unmedicated patients experience greater stress than medicated patients with OCD, in our study. A rodent quinpirole sensitization model of OCD showed a downregulation of Arc (a marker of plasticity-related neuronal activity) expressing neurons in the CA1 during stereotypical checking behaviour suggesting that the hippocampus may be involved in OCD more than is currently thought (Brozka, 2021).

There are some limitations to consider. First, even with automated segmentation, the small size of the amygdala poses challenges in accurately identifying its borders (Saygin et al., 2011). We also note that there is some evidence of poor test-retest reliability in segmentation of some hippocampal structures, including the medial, paralaminar nucleus, hippocampal fissure, and fimbria (Quattrini et al., 2020). Secondly, the cross-sectional nature of our study limits our interpretation of the effects of medication on subcortical volumes in OCD as these findings require validation using longitudinal studies. The third limitation is the inability of our study to account for heterogeneity in the clinical presentation of OCD in our models, particularly in light of published evidence for an association between OCD symptom profile and hippocampal volume (Reess et al., 2018). Lastly, due to lack of detailed information on medication status, we were unable to further investigate the effects of medication type, dosage, and duration on subfield volumes in medicated patients with OCD.

In summary, the association of medication status with volumetric alterations in OCD is consistent with previous work and emphasizes the importance of considering medication use as an important confounder in neuroimaging findings. Further investigation is required to elucidate the association between medication type, dosage, and duration and brain volumes in OCD over time.

## Acknowledgements

Supported by the Dutch Organization for Scientific Research (NWO) (grants 912-02-050, 907-00-012, 940-37-018, and 916.86.038); the Carlos III Health Institute (PI09/01331, PI10/01753, PI10/01003, CP10/00604, and CIBER-CB06/03/0034); the Agency for Administration of University and Research (AGAUR, Barcelona; 2009SGR1554); a “Miguel Servet” contract from the Carlos III Health Institute (CP10/00604) to Dr. Soriano-Mas; Ministry of Education, Culture, Sports, Science, and Technology (Japan) Grants-in-Aid for Young Scientists to Dr. Narumoto (23591724) and to Dr. Nakamae (24791223); Wellcome Trust project grant 064846; a grant from the Foundation for the Support of Research in the State of São Paulo (FAPESP) to Dr. Miguel (2005/55628-8); a FAPESP scholarship to Dr. Hoexter (2005/04206-6); and a National Research Foundation of Korea grant funded by the Korean government (Ministry of Education, Science, and Technology, 2012-0005150). High-performance computing (HPC) cluster at the University of Cape Town, South Africa.

## References

American Psychiatric Association. (2022). Diagnostic and statistical manual of mental disorders (5th ed., text rev.). https://doi.org/10.1176/appi.books.9780890425787

Atmaca, M., Ozdemir, H., Ozler, S., Kara, B., Ozler, Z., Kanmaz, E., Mermi, O., & Tezcan, E. (2008). Progress in Neuro-Psychopharmacology & Biological Psychiatry Hippocampus and amygdalar volumes in patients with refractory obsessive – compulsive disorder. 32, 1283–1286. https://doi.org/10.1016/j.pnpbp.2008.04.002

Barnes, J., Ridgway, G. R., Bartlett, J., Henley, S. M. D., Lehmann, M., Hobbs, N., Clarkson, M. J., MacManus, D. G., Ourselin, S., & Fox, N. C. (2010). Head size, age and gender adjustment in MRI studies: A necessary nuisance? NeuroImage, 53(4), 1244–1255. https://doi.org/10.1016/j.neuroimage.2010.06.025

Bartsch, T., Döhring, J., Reuter, S., Finke, C., Rohr, A., Brauer, H., Deuschl, G., & Jansen, O. (2015). Selective neuronal vulnerability of human hippocampal CA1 neurons: Lesion evolution, temporal course, and pattern of hippocampal damage in diffusion-weighted MR imaging. Journal of Cerebral Blood Flow and Metabolism, 35(11), 1836–1845. https://doi.org/10.1038/jcbfm.2015.137

Bernier, P. J., Bédard, A., Vinet, J., Lévesque, M., & Parent, A. (2002). Newly generated neurons in the amygdala and adjoining cortex of adult primates. Proceedings of the National Academy of Sciences of the United States of America, 99(17), 11464–11469. https://doi.org/10.1073/pnas.172403999

Boedhoe, P. S., Schmaal, L., Abe, Y., Ameis, S. H., Arnold, P. D., Batistuzzo, M. C., … & members of the ENIGMA OCD Working Group. (2017). Distinct subcortical volume alterations in pediatric and adult OCD: a worldwide meta-and mega-analysis. American Journal of Psychiatry, 174(1), 60–69.

Brown, E. M., Pierce, M. E., Clark, D. C., Fischl, B. R., Iglesias, J. E., Milberg, W. P., & Salat, D. H. (2020). Test-retest reliability of FreeSurfer automated hippocampal subfield segmentation within and across scanners. Neuroimage, 210, 116563.

Brožka, H. (2020). Hippocampus Dysfunction in Quinpirole Sensitization Model of Obsessive-Compulsive Disorder.

Brühl, A. B., Delsignore, A., Komossa, K., & Weidt, S. (2014). Neuroimaging in social anxiety disorder-A meta-analytic review resulting in a new neurofunctional model. Neuroscience and Biobehavioral Reviews, 47, 260–280. https://doi.org/10.1016/j.neubiorev.2014.08.003

Chen, L. W., Sun, D., Davis, S. L., Haswell, C. C., Dennis, E. L., Swanson, C. A., Whelan, C. D., Gutman, B., Jahanshad, N., Iglesias, J. E., Thompson, P., Wagner, H. R., Saemann, P., LaBar, K. S., & Morey, R. A. (2018). Smaller hippocampal CA1 subfield volume in posttraumatic stress disorder. Depression and Anxiety, 35(11), 1018–1029. https://doi.org/10.1002/da.22833

Cheng, D. T., Knight, D. C., Smith, C. N., Stein, E. A., & Helmstetter, F. J. (2003). Functional MRI of human amygdala activity during Pavlovian fear conditioning: Stimulus processing versus response expression. Behavioral Neuroscience, 117(1), 3–10. https://doi.org/10.1037/0735-7044.117.1.3

David, D. J., Samuels, B. A., Rainer, Q., Wang, J. W., Marsteller, D., Mendez, I., Drew, M., Craig, D. A., Guiard, B. P., Guilloux, J. P., Artymyshyn, R. P., Gardier, A. M., Gerald, C., Antonijevic, I. A., Leonardo, E. D., & Hen, R. (2009). Neurogenesis-Dependent and -Independent Effects of Fluoxetine in an Animal Model of Anxiety/Depression. Neuron, 62(4), 479–493. https://doi.org/10.1016/j.neuron.2009.04.017

De Wit, S. J., Alonso, P., Schweren, L., Mataix-Cols, D., Lochner, C., Menchón, J. M., Stein, D. J., Fouche, J. P., Soriano-Mas, C., Sato, J. R., Hoexter, M. Q., Denys, D., Nakamae, T., Nishida, S., Kwon, J. S., Jang, J. H., Busatto, G. F., Cardoner, N., Cath, D. C., … Van Den Heuvel, O. A. (2014). Multicenter voxel-based morphometry mega-analysis of structural brain scans in obsessive-compulsive disorder. American Journal of Psychiatry, 171(3), 340–349. https://doi.org/10.1176/appi.ajp.2013.13040574

Etkin, A. (2009). Functional neuroanatomy of anxiety: a neural circuit perspective. Behavioral neurobiology of anxiety and its treatment, 251–277.

Fischl, B., Salat, D. H., Busa, E., Albert, M., Dieterich, M., Haselgrove, C., Van Der Kouwe, A., Killiany, R., Kennedy, D., Klaveness, S., Montillo, A., Makris, N., Rosen, B., & Dale, A. M. (2002). Whole brain segmentation: Automated labeling of neuroanatomical structures in the human brain. Neuron, 33(3), 341–355. https://doi.org/10.1016/S0896-6273(02)00569-X

Fouche, J. P., Du Plessis, S., Hattingh, C., Roos, A., Lochner, C., Soriano-Mas, C., Sato, J. R., Nakamae, T., Nishida, S., Kwon, J. S., Jung, W. H., Mataix-Cols, D., Hoexter, M. Q., Alonso, P., De Wit, S. J., Veltman, D. J., Stein, D. J., & Van Den Heuvel, O. A. (2017). Cortical thickness in obsessive-compulsive disorder: Multisite mega-analysis of 780 brain scans from six centres. British Journal of Psychiatry, 210(1), 67–74. https://doi.org/10.1192/bjp.bp.115.164020

Fouche, J. P., Groenewold, N. A., Sevenoaks, T., Heany, S., Lochner, C., Alonso, P., … & Stein, D. J. (2022). Shape analysis of subcortical structures in obsessive-compulsive disorder and the relationship with comorbid anxiety, depression, and medication use: A meta-analysis by the OCD Brain Imaging Consortium. Brain and Behavior, 12(10), e2755.

Fowler, C. D., Liu, Y., Ouimet, C., & Wang, Z. (2002). The effects of social environment on adult neurogenesis in the female prairie vole. Journal of Neurobiology, 51(2), 115–128. https://doi.org/10.1002/neu.10042

González-García, I., & Visser, M. (2023, March). A Semantic Cognition Contribution to Mood and Anxiety Disorder Pathophysiology. In Healthcare (Vol. 11, No. 6, p. 821). MDPI.

Goodman, W. K., Price, L. H., Rasmussen, S. A., Mazure, C., Fleischmann, R. L., Hill, C. L., … & Charney, D. S. (1989). The Yale-Brown obsessive compulsive scale: I. Development, use, and reliability. Archives of general psychiatry, 46(11), 1006–1011.

Goodman, W. K., Grice, D. E., Lapidus, K. A. B., & Coffey, B. J. (2014). Obsessive-compulsive disorder. Psychiatric Clinics of North America, 37(3), 257–267. https://doi.org/10.1016/j.psc.2014.06.004

Green, S., & Vale, A. L. (1992). Role of amygdaloid nuclei in the anxiolytic effects of benzodiazepines in rats. Behavioural pharmacology, 3(3), 261–264.

Hodges, H., Green, S., & Glenn, B. (1987). Evidence that the amygdala is involved in benzodiazepine and serotonergic effects on punished responding but not on discrimination. Psychopharmacology, 92(4), 491–504. https://doi.org/10.1007/BF00176484

Iglesias, J. E., Augustinack, J. C., Nguyen, K., Player, C. M., Player, A., Wright, M., & Alzheimer’s Disease Neuroimaging Initiative. (2015). A computational atlas of the hippocampal formation using ex vivo, ultra-high resolution MRI: application to adaptive segmentation of in vivo MRI. Neuroimage, 115, 117–137.

Ivanov, I., Boedhoe, P. S. W., Abe, Y., Alonso, P., Ameis, S. H., Arnold, P. D., Balachander, S., Baker, J. T., Banaj, N., Bargalló, N., Batistuzzo, M. C., Benedetti, F., Beucke, J. C., Bollettini, I., Brem, S., Brennan, B. P., Buitelaar, J., Calvo, R., Cheng, Y., … Fouche, J.-P. (2022). Associations of medication with subcortical morphology across the lifespan in OCD: Results from the international ENIGMA Consortium. Journal of Affective Disorders, 318(August), 204–216. https://doi.org/10.1016/j.jad.2022.08.084

Kassem, M. S., Lagopoulos, J., Stait-Gardner, T., Price, W. S., Chohan, T. W., Arnold, J. C., Hatton, S. N., & Bennett, M. R. (2013). Stress-induced grey matter loss determined by MRI is primarily due to loss of dendrites and their synapses. Molecular Neurobiology, 47(2), 645–661. https://doi.org/10.1007/s12035-012-8365-7

Kwon, J. S., Shin, Y. W., Kim, C. W., Kim, Y. I., Youn, T., Han, M. H., … & Kim, J. J. (2003). Similarity and disparity of obsessive-compulsive disorder and schizophrenia in MR volumetric abnormalities of the hippocampus-amygdala complex. Journal of Neurology, Neurosurgery & Psychiatry, 74(7), 962–964.

LeDoux, J., 2007. The amygdala. Current biology, 17(20), pp.R868–R874.

Malberg, J. E., Eisch, A. J., Nestler, E. J., & Duman, R. S. (2000). Chronic antidepressant treatment increases neurogenesis in adult rat hippocampus. Journal of Neuroscience, 20(24), 9104–9110. https://doi.org/10.1523/jneurosci.20-24-09104.2000

Mandyam, C. D. (2013). The interplay between the hippocampus and amygdala in regulating aberrant hippocampal neurogenesis during protracted abstinence from alcohol dependence. Frontiers in Psychiatry, 4(JUN). https://doi.org/10.3389/fpsyt.2013.00061

Mataix-Cols, D., Do Rosario-Campos, M. C., & Leckman, J. F. (2005). A multidimensional model of obsessive-compulsive disorder. American Journal of Psychiatry, 162(2), 228–238. https://doi.org/10.1176/appi.ajp.162.2.228

McEwen, B. S. (1999). Stress and hippocampal plasticity. Annual Review of Neuroscience, 22, 105–122. https://doi.org/10.1146/annurev.neuro.22.1.105

Milad, M. R., Furtak, S. C., Greenberg, J. L., Keshaviah, A., Im, J. J., Falkenstein, M. J., Jenike, M., Rauch, S. L., & Wilhelm, S. (2013). Deficits in conditioned fear extinction in obsessive-compulsive disorder and neurobiological changes in the fear circuit. JAMA Psychiatry, 70(6), 608–618. https://doi.org/10.1001/jamapsychiatry.2013.914

Nacher, J., Lanuza, E., & McEwen, B. S. (2002). Distribution of PSA-NCAM expression in the amygdala of the adult rat. Neuroscience, 113(3), 479–484.

Nagy, J., Zámbó, K., & Decsi, L. (1979). Anti-anxiety action op diazepam after intraamygdaloid application in the rat. Neuropharmacology, 18(6), 573–576.

Nakagawa, S., & Cuthill, I. C. (2007). Effect size, confidence interval and statistical significance: A practical guide for biologists. Biological Reviews, 82(4), 591–605. https://doi.org/10.1111/j.1469-185X.2007.00027.x

Nazeer, A., Latif, F., Mondal, A., Azeem, M. W., & Greydanus, D. E. (2020). Obsessive-compulsive disorder in children and adolescents: epidemiology, diagnosis and management. Translational Pediatrics, 9, S76–S93. https://doi.org/10.21037/TP.2019.10.02

Nordenskjöld, R., Malmberg, F., Larsson, E. M., Simmons, A., Brooks, S. J., Lind, L., Ahlström, H., Johansson, L., & Kullberg, J. (2013). Intracranial volume estimated with commonly used methods could introduce bias in studies including brain volume measurements. NeuroImage, 83, 355–360. https://doi.org/10.1016/j.neuroimage.2013.06.068

Pauls, D. L., Abramovitch, A., Rauch, S. L., & Geller, D. A. (2014). Obsessive-compulsive disorder: An integrative genetic and neurobiological perspective. Nature Reviews Neuroscience, 15(6), 410–424. https://doi.org/10.1038/nrn3746

Perera, T. D., Coplan, J. D., Lisanby, S. H., Lipira, C. M., Arif, M., Carpio, C., Spitzer, G., Santarelli, L., Scharf, B., Hen, R., Rosoklija, G., Sackeim, H. A., & Dwork, A. J. (2007). Antidepressant-induced neurogenesis in the hippocampus of adult nonhuman primates. Journal of Neuroscience, 27(18), 4894–4901. https://doi.org/10.1523/JNEUROSCI.0237-07.2007

Quattrini, G., Pievani, M., Jovicich, J., Aiello, M., Bargalló, N., Barkhof, F., Bartres-Faz, D., Beltramello, A., Pizzini, F. B., Blin, O., Bordet, R., Caulo, M., Constantinides, M., Didic, M., Drevelegas, A., Ferretti, A., Fiedler, U., Floridi, P., Gros-Dagnac, H., … Marizzoni, M. (2020). Amygdalar nuclei and hippocampal subfields on MRI: Test-retest reliability of automated volumetry across different MRI sites and vendors. NeuroImage, 218(May). https://doi.org/10.1016/j.neuroimage.2020.116932

Rao, S., Raveendranathan, D., Shivakumar, V., Narayanaswamy, J. C., Venkatasubramanian, G., & Reddy, Y. J. (2018). Hippocampus volume alterations and the clinical correlates in medication naïve obsessive compulsive disorder. Journal of Affective Disorders, 236, 1–5.

Reess, T. J., Rus, O. G., Gürsel, D. A., Schmitz-Koep, B., Wagner, G., Berberich, G., & Koch, K. (2018). Association between hippocampus volume and symptom profiles in obsessive–compulsive disorder. NeuroImage: Clinical, 17(October 2017), 474–480. https://doi.org/10.1016/j.nicl.2017.11.006

Roeder, S. S., Burkardt, P., Rost, F., Rode, J., Brusch, L., Coras, R., … & Huttner, H. B. (2022). Evidence for postnatal neurogenesis in the human amygdala. Communications Biology, 5(1), 366.

Ruscio, A. M., Stein, D. J., Chiu, W. T., & Kessler, R. C. (2010). The epidemiology of obsessive-compulsive disorder in the National Comorbidity Survey Replication. Molecular Psychiatry, 15(1), 53–63. https://doi.org/10.1038/mp.2008.94

Ruscio, A.M., Stein, D.J., Chiu, W.T. and Kessler, R.C., 2010. The epidemiology of obsessive-compulsive disorder in the National Comorbidity Survey Replication. Molecular psychiatry, 15(1), p.53.

Sah, P., Faber, E. S. L., De Armentia, M. L., & Power, J. (2003). The amygdaloid complex: Anatomy and physiology. Physiological Reviews, 83(3), 803–834. https://doi.org/10.1152/physrev.00002.2003

Sämann, P. G., Iglesias, J. E., Gutman, B., Grotegerd, D., Leenings, R., Flint, C., Dannlowski, U., Clarke-Rubright, E. K., Morey, R. A., van Erp, T. G. M., Whelan, C. D., Han, L. K. M., van Velzen, L. S., Cao, B., Augustinack, J. C., Thompson, P. M., Jahanshad, N., & Schmaal, L. (2022). FreeSurfer-based segmentation of hippocampal subfields: A review of methods and applications, with a novel quality control procedure for ENIGMA studies and other collaborative efforts. Human Brain Mapping, 43(1), 207–233. https://doi.org/10.1002/hbm.25326

Santarelli, L., Saxe, M., Gross, C., Surget, A., Battaglia, F., Dulawa, S., Weisstaub, N., Lee, J., Duman, R., Arancio, O., Belzung, C., & Hen, R. (2003). Requirement of hippocampal neurogenesis for the behavioral effects of antidepressants. Science, 301(5634), 805–809. https://doi.org/10.1126/science.1083328

Sargolzaei, S., Sargolzaei, A., Cabrerizo, M., Chen, G., Goryawala, M., Noei, S., Zhou, Q., Duara, R., Barker, W., & Adjouadi, M. (2015). A practical guideline for intracranial volume estimation in patients with Alzheimer’s disease. BMC Bioinformatics, 16(7), 1–10. https://doi.org/10.1186/1471-2105-16-S7-S8

Saygin, Z. M., Kliemann, D., Iglesias, J. E., van der Kouwe, A. J. W., Boyd, E., Reuter, M., Stevens, A., Van Leemput, K., McKee, A., Frosch, M. P., Fischl, B., & Augustinack, J. C. (2017). High-resolution magnetic resonance imaging reveals nuclei of the human amygdala: manual segmentation to automatic atlas. NeuroImage, 155(April), 370–382. https://doi.org/10.1016/j.neuroimage.2017.04.046

Saygin, Z. M., Osher, D. E., Augustinack, J., Fischl, B., & Gabrieli, J. D. E. (2011). NeuroImage Connectivity-based segmentation of human amygdala nuclei using probabilistic tractography. NeuroImage, 56(3), 1353–1361. https://doi.org/10.1016/j.neuroimage.2011.03.006

Schmitz-Koep, B., Zimmermann, J., Menegaux, A., Nuttall, R., Bäuml, J.G., Schneider, S.C., Daamen, M., Boecker, H., Zimmer, C., Wolke, D. and Bartmann, P., 2021. Within amygdala: Basolateral parts are selectively impaired in premature-born adults. NeuroImage: Clinical, 31, p.102780.

Shapiro, L. A., Ng, K., Zhou, Q. Y., & Ribak, C. E. (2009). Subventricular zone-derived, newly generated neurons populate several olfactory and limbic forebrain regions. Epilepsy & Behavior, 14(1), 74–80.

Shi, H. J., Wang, S., Wang, X. P., Zhang, R. X., & Zhu, L. J. (2023). Hippocampus: Molecular, Cellular, and Circuit Features in Anxiety. Neuroscience Bulletin, 1–18.

Stein, D. J., Costa, D. L., Lochner, C., Miguel, E. C., Reddy, Y. C., Shavitt, R. G., & Simpson, H. B. (2019). Obsessive–compulsive disorder. Nature reviews Disease primers, 5(1), 1–21.

Tamminga, C. A., Stan, A. D., & Wagner, A. D. (2010). The hippocampal formation in schizophrenia. American Journal of Psychiatry, 167(10), 1178–1193.

Thorsen, A. L., Hagland, P., Radua, J., Mataix-Cols, D., Kvale, G., Hansen, B., & van den Heuvel, O. A. (2018). Emotional Processing in Obsessive-Compulsive Disorder: A Systematic Review and Meta-analysis of 25 Functional Neuroimaging Studies. Biological Psychiatry: Cognitive Neuroscience and Neuroimaging, 3(6), 563–571. https://doi.org/10.1016/j.bpsc.2018.01.009

van den Heuvel, O. A., Boedhoe, P. S. W., Bertolin, S., Bruin, W. B., Francks, C., Ivanov, I., Jahanshad, N., Kong, X. Z., Kwon, J. S., O’Neill, J., Paus, T., Patel, Y., Piras, F., Schmaal, L., Soriano-Mas, C., Spalletta, G., van Wingen, G. A., Yun, J. Y., Vriend, C., … Yamada, K. (2022). An overview of the first 5 years of the ENIGMA obsessive–compulsive disorder working group: The power of worldwide collaboration. Human Brain Mapping, 43(1), 23–36. https://doi.org/10.1002/hbm.24972

van den Heuvel, O. A., van Wingen, G., Soriano-Mas, C., Alonso, P., Chamberlain, S. R., Nakamae, T., Denys, D., Goudriaan, A. E., & Veltman, D. J. (2016). Brain circuitry of compulsivity. European Neuropsychopharmacology, 26(5), 810–827. https://doi.org/10.1016/j.euroneuro.2015.12.005

van den Heuvel, O.A., van Wingen, G., Soriano-Mas, C., Alonso, P., Chamberlain, S.R., Nakamae, T., Denys, D., Goudriaan, A.E. and Veltman, D.J., 2016. Brain circuitry of compulsivity. European Neuropsychopharmacology, 26(5), pp.810–827.

van der Meer, D., Rokicki, J., Kaufmann, T., Córdova-Palomera, A., Moberget, T., Alnæs, D., Bettella, F., Frei, O., Doan, N. T., Sønderby, I. E., Smeland, O. B., Agartz, I., Bertolino, A., Bralten, J., Brandt, C. L., Buitelaar, J. K., Djurovic, S., van Donkelaar, M., Dørum, E. S., … Westlye, L. T. (2018). Brain scans from 21,297 individuals reveal the genetic architecture of hippocampal subfield volumes. Molecular Psychiatry. https://doi.org/10.1038/s41380-018-0262-7

van Oudheusden, L. J. B., van de Schoot, R., Hoogendoorn, A., van Oppen, P., Kaarsemaker, M., Meynen, G., & van Balkom, A. J. L. M. (2020). Classification of comorbidity in obsessive–compulsive disorder: A latent class analysis. Brain and Behavior, 10(7), 1–10. https://doi.org/10.1002/brb3.1641

Vattimo, E. F. Q., dos Santos, A. C., Hoexter, M. Q., Frudit, P., Miguel, E. C., Shavitt, R. G., & Batistuzzo, M. C. (2021). Higher volumes of hippocampal subfields in pediatric obsessive-compulsive disorder. Psychiatry Research - Neuroimaging, 307(September), 111200. https://doi.org/10.1016/j.pscychresns.2020.111200

Wang, J. W., David, D. J., Monckton, J. E., Battaglia, F., & Hen, R. (2008). Chronic fluoxetine stimulates maturation and synaptic plasticity of adult-born hippocampal granule cells. Journal of Neuroscience, 28(6), 1374–1384. https://doi.org/10.1523/JNEUROSCI.3632-07.2008

Weeland, C. J., Vriend, C., van der Werf, Y., Huyser, C., Hillegers, M., Tiemeier, H.,& van den Heuvel, O. A. (2022). Thalamic subregions and obsessive-compulsive symptoms in 2,500 children from the general population. Journal of the American Academy of Child & Adolescent Psychiatry, 61(2), 321–330.

Zangrossi Jr, H., Viana, M.B. and Graeff, F.G., 1999. Anxiolytic effect of intra-amygdala injection of midazolam and 8-hydroxy-2-(di-n-propylamino) tetralin in the elevated T-maze. European Journal of Pharmacology, 369(3), pp.267–270.

Zhang, L., Hu, X., Lu, L., Li, B., Hu, X., Bu, X., Li, H., Tang, S., Yang, Y., Roberts, N., Sweeney, J. A., Gong, Q., & Huang, X. (2019). Abnormalities of hippocampal shape and subfield volumes in medication-free patients with obsessive–compulsive disorder. Human Brain Mapping, 40(14), 4105–4113. https://doi.org/10.1002/hbm.24688

Zhang, L., Hu, X., Lu, L., Li, B., Hu, X., Bu, X., & Huang, X. (2020). Anatomic alterations across amygdala subnuclei in medication-free patients with obsessive–compulsive disorder. Journal of Psychiatry and Neuroscience, 45(5), 334–343.

