## Supplemental data for "Hippocampal and amygdala subfield volumes in obsessive-compulsive disorder differ according to medication status"

**Supplemental table S1a:** Mega analysis linear regression results for **bilateral** amygdala and hippocampal subnuclei volumes for OCD patients (n=381) compared with adult healthy controls (HCs) (n=338), adjusting for age, sex and scan site and whole volumes for amygdala and hippocampus.

|  | **Cohen’s d**  **(OCD vs HCs)** | **Standard error of *d*** | **FDR corrected**  **p-value** | **Uncorrected p-value** |
| --- | --- | --- | --- | --- |
| Lateral | 0.095 | 0.075 | 0.353 | 0.168 |
| Basal | 0.102 | 0.075 | 0.353 | 0.149 |
| Accessory Basal | 0.017 | 0.075 | 0.935 | 0.802 |
| AAA | -0.050 | 0.075 | 0.664 | 0.419 |
| Central | 0.036 | 0.075 | 0.828 | 0.631 |
| Medial | -0.076 | 0.075 | 0.564 | 0.296 |
| Cortical | 0.004 | 0.075 | 0.979 | 0.956 |
| Corticoamygdaloid transition | 0.002 | 0.075 | 0.979 | 0.979 |
| Paralaminar | 0.154 | 0.075 | 0.150 | 0.029 |
| Parasubiculum | 0.051 | 0.075 | 0.664 | 0.443 |
| Presibiculum | 0.152 | 0.075 | 0.156 | 0.037 |
| Subiculum | 0.039 | 0.075 | 0.756 | 0.540 |
| CA1 | -0.196 | 0.075 | 0.063 | 0.006 |
| CA3 | -0.096 | 0.075 | 0.353 | 0.164 |
| CA4 | -0.111 | 0.075 | 0.353 | 0.104 |
| GC ML DG | -0.155 | 0.075 | 0.150 | 0.023 |
| Molecular layer | -0.207 | 0.075 | 0.060 | 0.003 |
| HATA | -0.049 | 0.075 | 0.636 | 0.363 |
| Fimbria | 0.108 | 0.075 | 0.353 | 0.122 |
| Hippocampal fissure | 0.012 | 0.075 | 0.955 | 0.864 |
| Hippocampal tail | 0.030 | 0.075 | 0.841 | 0.681 |

Abbreviations: cornu ammonis (CA) sectors, CA1, CA2-3, CA4, granule cell layer of dentate gyrus (DG), molecular layer (ML), hippocampus–amygdala transition area (HATA), corticoamygdaloid transition area (CAT), anterior amygdaloid area (AAA).

**Supplemental table S1b:** Mega analysis linear regression results for **separate left and right** amygdala and hippocampal subnuclei volumes for OCD patients (n=381) compared with adult healthy controls (HCs) (n=338), adjusting for age, sex and scan site and whole volumes for amygdala and hippocampus.

|  | **Cohen’s d**  **(OCD vs HCs)** | **Standard error of *d*** | **FDR corrected**  **p-value** | **Uncorrected p-value** |
| --- | --- | --- | --- | --- |
| L Lateral | 0.077 | 0.075 | 0.514 | 0.245 |
| L Basal | 0.094 | 0.075 | 0.473 | 0.185 |
| L Accessory Basal | 0.022 | 0.075 | 0.813 | 0.755 |
| L AAA | -0.052 | 0.075 | 0.701 | 0.417 |
| L Central | 0.099 | 0.075 | 0.473 | 0.188 |
| L Medial | -0.050 | 0.075 | 0.706 | 0.500 |
| L Cortical | 0.034 | 0.075 | 0.738 | 0.633 |
| L Corticoamygdaloid transition | -0.044 | 0.075 | 0.706 | 0.521 |
| L Paralaminar | 0.111 | 0.075 | 0.389 | 0.120 |
| R Lateral | 0.095 | 0.075 | 0.473 | 0.191 |
| R Basal | 0.086 | 0.075 | 0.514 | 0.231 |
| R Accessory Basal | 0.003 | 0.075 | 0.972 | 0.962 |
| R AAA | -0.038 | 0.075 | 0.725 | 0.552 |
| R Central | -0.042 | 0.075 | 0.730 | 0.574 |
| R Medial | -0.086 | 0.075 | 0.514 | 0.240 |
| R Cortical | -0.035 | 0.075 | 0.738 | 0.623 |
| R Corticoamygdaloid transition | 0.044 | 0.075 | 0.702 | 0.456 |
| R Paralaminar | 0.161 | 0.075 | 0.194 | 0.023 |
| L Parasubiculum | -0.002 | 0.075 | 0.972 | 0.972 |
| L Presubiculum | 0.134 | 0.075 | 0.322 | 0.069 |
| L Subiculum | 0.027 | 0.075 | 0.763 | 0.690 |
| L CA1 | -0.149 | 0.075 | 0.223 | 0.033 |
| L CA3 | -0.064 | 0.075 | 0.642 | 0.367 |
| L CA4 | -0.077 | 0.075 | 0.556 | 0.278 |
| L GC ML DG | -0.110 | 0.075 | 0.389 | 0.115 |
| L Molecular layer | -0.175 | 0.075 | 0.170 | 0.013 |
| L HATA | -0.064 | 0.075 | 0.565 | 0.296 |
| L Fimbria | 0.052 | 0.075 | 0.702 | 0.468 |
| L Hippocampal fissure | -0.046 | 0.075 | 0.706 | 0.521 |
| L Hippocampal tail | 0.020 | 0.075 | 0.831 | 0.791 |
| R Parasubiculum | 0.095 | 0.075 | 0.460 | 0.153 |
| R Presubiculum | 0.145 | 0.075 | 0.250 | 0.048 |
| R Subiculum | 0.047 | 0.075 | 0.702 | 0.451 |
| R CA1 | -0.193 | 0.075 | 0.170 | 0.009 |
| R CA3 | -0.111 | 0.075 | 0.389 | 0.109 |
| R CA4 | -0.120 | 0.075 | 0.324 | 0.077 |
| R Molecular layer | -0.190 | 0.075 | 0.170 | 0.006 |
| R GC ML DG | -0.166 | 0.075 | 0.170 | 0.016 |
| R HATA | -0.025 | 0.075 | 0.746 | 0.658 |
| R Fimbria | 0.144 | 0.075 | 0.223 | 0.037 |
| R Hippocampal fissure | 0.065 | 0.075 | 0.642 | 0.358 |
| R Hippocampal tail | 0.035 | 0.075 | 0.738 | 0.627 |

Abbreviations: cornu ammonis (CA) sectors, CA1, CA2-3, CA4, granule cell layer of dentate gyrus (DG), molecular layer (ML), hippocampus–amygdala transition area (HATA), corticoamygdaloid transition area (CAT), anterior amygdaloid area (AAA).

**Supplemental table S2a:** Mega analysis linear regression results for **bilateral** amygdala and hippocampal subnuclei volumes for OCD patients (n=369) **without PTSD** compared with adult healthy controls (HCs) (n=338), adjusting for age, sex and scan site and whole volumes for amygdala and hippocampus.

|  | **Cohen’s d**  **(OCD vs HCs)** | **Standard error of *d*** | **FDR corrected**  **p-value** | **Uncorrected p-value** |
| --- | --- | --- | --- | --- |
| Lateral | 0.094 | 0.075 | 0.371 | 0.177 |
| Basal | 0.115 | 0.075 | 0.318 | 0.106 |
| Accessory Basal | 0.016 | 0.075 | 0.903 | 0.817 |
| AAA | -0.048 | 0.075 | 0.673 | 0.442 |
| Central | 0.051 | 0.075 | 0.673 | 0.495 |
| Medial | -0.078 | 0.075 | 0.546 | 0.286 |
| Cortical | -0.010 | 0.075 | 0.922 | 0.891 |
| Corticoamygdaloid transition | -0.006 | 0.075 | 0.922 | 0.922 |
| Paralaminar | 0.156 | 0.075 | 0.142 | 0.028 |
| Parasubiculum | 0.044 | 0.075 | 0.673 | 0.512 |
| Presibiculum | 0.156 | 0.075 | 0.142 | 0.034 |
| Subiculum | 0.046 | 0.075 | 0.673 | 0.474 |
| CA1 | -0.201 | 0.076 | 0.054 | 0.005 |
| CA3 | -0.097 | 0.075 | 0.371 | 0.164 |
| CA4 | -0.120 | 0.075 | 0.288 | 0.082 |
| GC ML DG | -0.166 | 0.075 | 0.111 | 0.016 |
| **Molecular layer** | **-0.214** | **0.076** | **0.045** | **0.002** |
| HATA | -0.048 | 0.075 | 0.655 | 0.374 |
| Fimbria | 0.107 | 0.075 | 0.336 | 0.128 |
| Hippocampal fissure | 0.029 | 0.075 | 0.794 | 0.680 |
| Hippocampal tail | 0.034 | 0.075 | 0.794 | 0.648 |

Abbreviations: cornu ammonis (CA) sectors, CA1, CA2-3, CA4, granule cell layer of dentate gyrus (DG), molecular layer (ML), hippocampus–amygdala transition area (HATA), corticoamygdaloid transition area (CAT), anterior amygdaloid area (AAA).

**Supplemental table S2b:** Mega analysis linear regression results **for left and right** amygdala and hippocampal subnuclei volumes for OCD patients (n=369) **without PTSD** compared with adult healthy controls (HCs) (n=338), adjusting for age, sex and scan site and whole volumes for amygdala and hippocampus.

|  | **Cohen’s d**  **(OCD vs HCs)** | **Standard error of *d*** | **FDR corrected**  **p-value** | **Uncorrected p-value** |
| --- | --- | --- | --- | --- |
| L Lateral | 0.079 | 0.075 | 0.519 | 0.241 |
| L Basal | 0.109 | 0.075 | 0.386 | 0.129 |
| L Accessory Basal | 0.019 | 0.075 | 0.860 | 0.798 |
| L AAA | -0.052 | 0.075 | 0.659 | 0.423 |
| L Central | 0.106 | 0.075 | 0.455 | 0.162 |
| L Medial | -0.056 | 0.075 | 0.659 | 0.450 |
| L Cortical | 0.016 | 0.075 | 0.863 | 0.822 |
| L Corticoamygdaloid transition | -0.052 | 0.075 | 0.659 | 0.455 |
| L Paralaminar | 0.116 | 0.075 | 0.344 | 0.106 |
| R Lateral | 0.091 | 0.075 | 0.495 | 0.212 |
| R Basal | 0.095 | 0.075 | 0.464 | 0.188 |
| R Accessory Basal | 0.005 | 0.075 | 0.944 | 0.944 |
| R AAA | -0.034 | 0.075 | 0.763 | 0.599 |
| R Central | -0.023 | 0.075 | 0.856 | 0.758 |
| R Medial | -0.083 | 0.075 | 0.521 | 0.261 |
| R Cortical | -0.039 | 0.075 | 0.763 | 0.586 |
| R Corticoamygdaloid transition | 0.038 | 0.075 | 0.738 | 0.527 |
| R Paralaminar | 0.159 | 0.075 | 0.218 | 0.026 |
| L Parasubiculum | -0.010 | 0.075 | 0.907 | 0.886 |
| L Presubiculum | 0.138 | 0.075 | 0.265 | 0.063 |
| L Subiculum | 0.031 | 0.075 | 0.800 | 0.653 |
| L CA1 | -0.151 | 0.075 | 0.218 | 0.031 |
| L CA3 | -0.063 | 0.075 | 0.635 | 0.378 |
| L CA4 | -0.083 | 0.075 | 0.519 | 0.247 |
| L GC ML DG | -0.119 | 0.075 | 0.344 | 0.092 |
| L Molecular layer | -0.184 | 0.075 | 0.120 | 0.010 |
| L HATA | -0.063 | 0.075 | 0.557 | 0.305 |
| L Fimbria | 0.055 | 0.075 | 0.659 | 0.450 |
| L Hippocampal fissure | -0.028 | 0.075 | 0.812 | 0.696 |
| L Hippocampal tail | 0.022 | 0.075 | 0.856 | 0.774 |
| R Parasubiculum | 0.090 | 0.075 | 0.464 | 0.178 |
| R Presubiculum | 0.147 | 0.075 | 0.240 | 0.046 |
| R Subiculum | 0.056 | 0.075 | 0.635 | 0.376 |
| R CA1 | -0.199 | 0.075 | 0.120 | 0.007 |
| R CA3 | -0.113 | 0.075 | 0.344 | 0.105 |
| R CA4 | -0.129 | 0.075 | 0.265 | 0.059 |
| R Molecular layer | -0.193 | 0.075 | 0.120 | 0.005 |
| R GC ML DG | -0.176 | 0.075 | 0.120 | 0.011 |
| R HATA | -0.024 | 0.075 | 0.800 | 0.666 |
| R Fimbria | 0.139 | 0.075 | 0.240 | 0.045 |
| R Hippocampal fissure | 0.076 | 0.075 | 0.529 | 0.277 |
| R Hippocampal tail | 0.039 | 0.075 | 0.763 | 0.591 |

Abbreviations: cornu ammonis (CA) sectors, CA1, CA2-3, CA4, granule cell layer of dentate gyrus (DG), molecular layer (ML), hippocampus–amygdala transition area (HATA), corticoamygdaloid transition area (CAT), anterior amygdaloid area (AAA).

**Supplemental table S3a:** Mega analysis linear regression results for **bilateral** amygdala and hippocampal subnuclei volumes for OCD patients (n=74) **with anxiety disorder comorbidity** compared with adult healthy controls (HCs) (n=338), adjusting for age, sex and scan site and whole volumes for amygdala and hippocampus.

|  | **Cohen’s d**  **(OCD vs HCs)** | **Standard error of *d*** | **FDR corrected**  **p-value** | **Uncorrected p-value** |
| --- | --- | --- | --- | --- |
| Lateral | 0.010 | 0.099 | 0.980 | 0.933 |
| Basal | 0.082 | 0.099 | 0.800 | 0.495 |
| Accessory Basal | 0.146 | 0.099 | 0.491 | 0.234 |
| AAA | -0.052 | 0.099 | 0.835 | 0.634 |
| Central | 0.111 | 0.099 | 0.684 | 0.391 |
| Medial | -0.053 | 0.099 | 0.835 | 0.676 |
| Cortical | 0.212 | 0.099 | 0.319 | 0.076 |
| Corticoamygdaloid transition | 0.147 | 0.099 | 0.433 | 0.186 |
| Paralaminar | 0.115 | 0.099 | 0.670 | 0.351 |
| Parasubiculum | 0.051 | 0.099 | 0.835 | 0.657 |
| Presibiculum | 0.230 | 0.099 | 0.319 | 0.063 |
| Subiculum | 0.053 | 0.099 | 0.835 | 0.629 |
| CA1 | -0.231 | 0.099 | 0.319 | 0.066 |
| CA3 | -0.170 | 0.099 | 0.419 | 0.160 |
| CA4 | -0.167 | 0.099 | 0.419 | 0.156 |
| GC ML DG | -0.182 | 0.099 | 0.419 | 0.125 |
| Molecular layer | -0.253 | 0.099 | 0.319 | 0.036 |
| HATA | -0.013 | 0.099 | 0.980 | 0.891 |
| Fimbria | 0.223 | 0.099 | 0.319 | 0.064 |
| Hippocampal fissure | -0.035 | 0.099 | 0.906 | 0.777 |
| Hippocampal tail | 0.001 | 0.099 | 0.993 | 0.993 |

**Supplemental table S3b:** Mega analysis linear regression results for **left and right** amygdala and hippocampal subnuclei volumes for OCD patients (n=74) **with anxiety disorder comorbidity** compared with adult healthy controls (HCs) (n=338), adjusting for age, sex and scan site and whole volumes for amygdala and hippocampus.

|  | **Cohen’s d**  **(OCD vs HCs)** | **Standard error of *d*** | **FDR corrected**  **p-value** | **Uncorrected p-value** |
| --- | --- | --- | --- | --- |
| L Lateral | -0.009 | 0.099 | 0.974 | 0.938 |
| L Basal | 0.077 | 0.099 | 0.796 | 0.531 |
| L Accessory Basal | 0.087 | 0.099 | 0.760 | 0.489 |
| L AAA | -0.079 | 0.099 | 0.760 | 0.487 |
| L Central | 0.038 | 0.099 | 0.974 | 0.769 |
| L Medial | -0.092 | 0.099 | 0.760 | 0.471 |
| L Cortical | 0.183 | 0.099 | 0.563 | 0.139 |
| L Corticoamygdaloid transition | 0.105 | 0.099 | 0.756 | 0.383 |
| L Paralaminar | 0.096 | 0.099 | 0.760 | 0.443 |
| R Lateral | 0.037 | 0.099 | 0.974 | 0.766 |
| R Basal | 0.058 | 0.099 | 0.888 | 0.634 |
| R Accessory Basal | 0.172 | 0.099 | 0.563 | 0.160 |
| R AAA | -0.017 | 0.099 | 0.974 | 0.880 |
| R Central | 0.147 | 0.099 | 0.604 | 0.259 |
| R Medial | -0.031 | 0.099 | 0.974 | 0.811 |
| R Cortical | 0.167 | 0.099 | 0.563 | 0.173 |
| R Corticoamygdaloid transition | 0.162 | 0.099 | 0.563 | 0.123 |
| R Paralaminar | 0.104 | 0.099 | 0.756 | 0.407 |
| L Parasubiculum | 0.010 | 0.099 | 0.974 | 0.936 |
| L Presubiculum | 0.259 | 0.099 | 0.563 | 0.041 |
| L Subiculum | 0.111 | 0.099 | 0.734 | 0.350 |
| L CA1 | -0.226 | 0.099 | 0.563 | 0.064 |
| L CA3 | -0.151 | 0.099 | 0.568 | 0.230 |
| L CA4 | -0.181 | 0.099 | 0.563 | 0.142 |
| L GC ML DG | -0.174 | 0.099 | 0.563 | 0.154 |
| L Molecular layer | -0.223 | 0.099 | 0.563 | 0.071 |
| L HATA | 0.004 | 0.099 | 0.974 | 0.974 |
| L Fimbria | 0.155 | 0.099 | 0.563 | 0.214 |
| L Hippocampal fissure | -0.063 | 0.099 | 0.888 | 0.619 |
| L Hippocampal tail | 0.035 | 0.099 | 0.974 | 0.785 |
| R Parasubiculum | 0.095 | 0.099 | 0.756 | 0.414 |
| R Presubiculum | 0.157 | 0.099 | 0.563 | 0.207 |
| R Subiculum | -0.006 | 0.099 | 0.974 | 0.958 |
| R CA1 | -0.163 | 0.099 | 0.563 | 0.208 |
| R CA3 | -0.157 | 0.099 | 0.563 | 0.187 |
| R CA4 | -0.122 | 0.099 | 0.646 | 0.292 |
| R Molecular layer | -0.214 | 0.099 | 0.563 | 0.071 |
| R GC ML DG | -0.150 | 0.099 | 0.563 | 0.212 |
| R HATA | -0.015 | 0.099 | 0.974 | 0.877 |
| R Fimbria | 0.260 | 0.099 | 0.563 | 0.031 |
| R Hippocampal fissure | 0.007 | 0.099 | 0.974 | 0.953 |
| R Hippocampal tail | -0.016 | 0.099 | 0.974 | 0.897 |

Abbreviations: cornu ammonis (CA) sectors, CA1, CA2-3, CA4, granule cell layer of dentate gyrus (DG), molecular layer (ML), hippocampus–amygdala transition area (HATA), corticoamygdaloid transition area (CAT), anterior amygdaloid area (AAA).

**Supplemental table S4a:** Mega analysis linear regression results for **bilateral** amygdala and hippocampal subnuclei volumes for OCD patients (n=307) **without anxiety disorder comorbidity** compared with adult healthy controls (HCs) (n=338), adjusting for age, sex and scan site and whole volumes for amygdala and hippocampus.

|  | **Cohen’s d**  **(OCD vs HCs)** | **Standard error of *d*** | **FDR corrected**  **p-value** | **Uncorrected p-value** |
| --- | --- | --- | --- | --- |
| Lateral | 0.108 | 0.079 | 0.435 | 0.135 |
| Basal | 0.105 | 0.079 | 0.435 | 0.161 |
| Accessory Basal | -0.026 | 0.079 | 0.777 | 0.728 |
| AAA | -0.054 | 0.079 | 0.568 | 0.406 |
| Central | 0.026 | 0.079 | 0.777 | 0.741 |
| Medial | -0.086 | 0.079 | 0.501 | 0.263 |
| Cortical | -0.062 | 0.079 | 0.568 | 0.393 |
| Corticoamygdaloid transition | -0.042 | 0.079 | 0.666 | 0.530 |
| Paralaminar | 0.156 | 0.079 | 0.188 | 0.036 |
| Parasubiculum | 0.064 | 0.079 | 0.568 | 0.354 |
| Presibiculum | 0.150 | 0.079 | 0.222 | 0.053 |
| Subiculum | 0.038 | 0.079 | 0.666 | 0.571 |
| CA1 | -0.188 | 0.079 | 0.129 | 0.012 |
| CA3 | -0.084 | 0.079 | 0.501 | 0.247 |
| CA4 | -0.099 | 0.079 | 0.435 | 0.166 |
| GC ML DG | -0.153 | 0.079 | 0.188 | 0.033 |
| Molecular layer | -0.203 | 0.079 | 0.117 | 0.006 |
| HATA | -0.058 | 0.079 | 0.546 | 0.312 |
| Fimbria | 0.089 | 0.079 | 0.501 | 0.224 |
| Hippocampal fissure | 0.021 | 0.079 | 0.777 | 0.777 |
| Hippocampal tail | 0.046 | 0.079 | 0.666 | 0.553 |

Abbreviations: cornu ammonis (CA) sectors, CA1, CA2-3, CA4, granule cell layer of dentate gyrus (DG), molecular layer (ML), hippocampus–amygdala transition area (HATA), corticoamygdaloid transition area (CAT), anterior amygdaloid area (AAA).

**Supplemental table S4b:** Mega analysis linear regression results for **left and right** amygdala and hippocampal subnuclei volumes for OCD patients (n=307) **without anxiety disorder comorbidity** compared with adult healthy controls (HCs) (n=338), adjusting for age, sex and scan site and whole volumes for amygdala and hippocampus.

|  | **Cohen’s d**  **(OCD vs HCs)** | **Standard error of *d*** | **FDR corrected**  **p-value** | **Uncorrected p-value** |
| --- | --- | --- | --- | --- |
| L Lateral | 0.103 | 0.079 | 0.398 | 0.144 |
| L Basal | 0.102 | 0.079 | 0.398 | 0.173 |
| L Accessory Basal | 0.000 | 0.079 | 0.996 | 0.996 |
| L AAA | -0.045 | 0.079 | 0.677 | 0.506 |
| L Central | 0.115 | 0.079 | 0.398 | 0.147 |
| L Medial | -0.044 | 0.079 | 0.688 | 0.574 |
| L Cortical | -0.015 | 0.079 | 0.934 | 0.845 |
| L Corticoamygdaloid transition | -0.085 | 0.079 | 0.490 | 0.245 |
| L Paralaminar | 0.110 | 0.079 | 0.398 | 0.145 |
| R Lateral | 0.094 | 0.079 | 0.458 | 0.218 |
| R Basal | 0.082 | 0.079 | 0.525 | 0.281 |
| R Accessory Basal | -0.055 | 0.079 | 0.650 | 0.455 |
| R AAA | -0.052 | 0.079 | 0.650 | 0.441 |
| R Central | -0.072 | 0.079 | 0.574 | 0.355 |
| R Medial | -0.109 | 0.079 | 0.398 | 0.160 |
| R Cortical | -0.100 | 0.079 | 0.398 | 0.180 |
| R Corticoamygdaloid transition | 0.005 | 0.079 | 0.963 | 0.940 |
| R Paralaminar | 0.165 | 0.079 | 0.231 | 0.027 |
| L Parasubiculum | 0.007 | 0.079 | 0.963 | 0.919 |
| L Presubiculum | 0.119 | 0.079 | 0.398 | 0.127 |
| L Subiculum | 0.007 | 0.079 | 0.963 | 0.922 |
| L CA1 | -0.136 | 0.079 | 0.356 | 0.063 |
| L CA3 | -0.048 | 0.079 | 0.677 | 0.516 |
| L CA4 | -0.060 | 0.079 | 0.650 | 0.424 |
| L GC ML DG | -0.103 | 0.079 | 0.398 | 0.163 |
| L Molecular layer | -0.179 | 0.079 | 0.219 | 0.016 |
| L HATA | -0.068 | 0.079 | 0.525 | 0.295 |
| L Fimbria | 0.030 | 0.079 | 0.814 | 0.698 |
| L Hippocampal fissure | -0.043 | 0.079 | 0.688 | 0.567 |
| L Hippocampal tail | 0.024 | 0.079 | 0.865 | 0.762 |
| R Parasubiculum | 0.107 | 0.079 | 0.398 | 0.123 |
| R Presubiculum | 0.158 | 0.079 | 0.293 | 0.042 |
| R Subiculum | 0.065 | 0.079 | 0.542 | 0.322 |
| R CA1 | -0.194 | 0.079 | 0.219 | 0.013 |
| R CA3 | -0.106 | 0.079 | 0.398 | 0.148 |
| R CA4 | -0.116 | 0.079 | 0.398 | 0.104 |
| R Molecular layer | -0.176 | 0.079 | 0.219 | 0.015 |
| R GC ML DG | -0.169 | 0.079 | 0.219 | 0.021 |
| R HATA | -0.037 | 0.079 | 0.680 | 0.534 |
| R Fimbria | 0.133 | 0.079 | 0.356 | 0.068 |
| R Hippocampal fissure | 0.077 | 0.079 | 0.525 | 0.300 |
| R Hippocampal tail | 0.056 | 0.079 | 0.650 | 0.464 |

Abbreviations: cornu ammonis (CA) sectors, CA1, CA2-3, CA4, granule cell layer of dentate gyrus (DG), molecular layer (ML), hippocampus–amygdala transition area (HATA), corticoamygdaloid transition area (CAT), anterior amygdaloid area (AAA).

**Supplemental table S5a:** Mega analysis linear regression results for **bilateral** amygdala and hippocampal subnuclei volumes for OCD patients (n=95) **with MDD comorbidity** compared with adult healthy controls (HCs) (n=338), adjusting for age, sex and scan site and whole volumes for amygdala and hippocampus.

|  | **Cohen’s d**  **(OCD vs HCs)** | **Standard error of *d*** | **FDR corrected**  **p-value** | **Uncorrected p-value** |
| --- | --- | --- | --- | --- |
| Lateral | 0.123 | 0.096 | 0.747 | 0.249 |
| Basal | 0.023 | 0.096 | 0.888 | 0.831 |
| Accessory Basal | -0.032 | 0.096 | 0.888 | 0.771 |
| AAA | -0.032 | 0.096 | 0.888 | 0.742 |
| Central | -0.042 | 0.096 | 0.888 | 0.723 |
| Medial | -0.045 | 0.096 | 0.888 | 0.690 |
| Cortical | 0.032 | 0.096 | 0.888 | 0.767 |
| Corticoamygdaloid transition | 0.014 | 0.096 | 0.888 | 0.887 |
| Paralaminar | 0.135 | 0.096 | 0.747 | 0.224 |
| Parasubiculum | 0.024 | 0.096 | 0.888 | 0.817 |
| Presibiculum | 0.178 | 0.097 | 0.612 | 0.117 |
| Subiculum | -0.014 | 0.096 | 0.888 | 0.888 |
| CA1 | -0.279 | 0.097 | 0.144 | 0.014 |
| CA3 | -0.105 | 0.096 | 0.748 | 0.334 |
| CA4 | -0.098 | 0.096 | 0.748 | 0.356 |
| GC ML DG | -0.140 | 0.096 | 0.747 | 0.191 |
| Molecular layer | -0.327 | 0.097 | 0.053 | 0.003 |
| HATA | -0.025 | 0.096 | 0.888 | 0.771 |
| Fimbria | 0.192 | 0.097 | 0.527 | 0.075 |
| Hippocampal fissure | -0.036 | 0.096 | 0.888 | 0.744 |
| Hippocampal tail | 0.109 | 0.096 | 0.748 | 0.340 |

Abbreviations: cornu ammonis (CA) sectors, CA1, CA2-3, CA4, granule cell layer of dentate gyrus (DG), molecular layer (ML), hippocampus–amygdala transition area (HATA), corticoamygdaloid transition area (CAT), anterior amygdaloid area (AAA).

**Supplemental table S5b:** Mega analysis linear regression results for **left and right** amygdala and hippocampal subnuclei volumes for OCD patients (n=95) **with MDD comorbidity** compared with adult healthy controls (HCs) (n=338), adjusting for age, sex and scan site and whole volumes for amygdala and hippocampus.

|  | **Cohen’s d**  **(OCD vs HCs)** | **Standard error of *d*** | **FDR corrected**  **p-value** | **Uncorrected p-value** |
| --- | --- | --- | --- | --- |
| L Lateral | 0.038 | 0.096 | 0.975 | 0.711 |
| L Basal | 0.021 | 0.096 | 0.975 | 0.848 |
| L Accessory Basal | -0.001 | 0.096 | 0.996 | 0.996 |
| L AAA | -0.102 | 0.096 | 0.903 | 0.306 |
| L Central | -0.053 | 0.096 | 0.975 | 0.651 |
| L Medial | -0.061 | 0.096 | 0.975 | 0.596 |
| L Cortical | 0.019 | 0.096 | 0.975 | 0.863 |
| L Corticoamygdaloid transition | 0.024 | 0.096 | 0.975 | 0.827 |
| L Paralaminar | 0.130 | 0.096 | 0.858 | 0.245 |
| R Lateral | 0.193 | 0.097 | 0.555 | 0.087 |
| R Basal | 0.020 | 0.096 | 0.975 | 0.856 |
| R Accessory Basal | -0.059 | 0.096 | 0.975 | 0.591 |
| R AAA | 0.047 | 0.096 | 0.975 | 0.639 |
| R Central | -0.032 | 0.096 | 0.975 | 0.781 |
| R Medial | -0.027 | 0.096 | 0.975 | 0.814 |
| R Cortical | 0.033 | 0.096 | 0.975 | 0.766 |
| R Corticoamygdaloid transition | 0.003 | 0.096 | 0.996 | 0.974 |
| R Paralaminar | 0.106 | 0.096 | 0.903 | 0.344 |
| L Parasubiculum | 0.016 | 0.096 | 0.975 | 0.883 |
| L Presubiculum | 0.265 | 0.097 | 0.231 | 0.022 |
| L Subiculum | 0.022 | 0.096 | 0.975 | 0.834 |
| L CA1 | -0.185 | 0.097 | 0.555 | 0.092 |
| L CA3 | -0.137 | 0.096 | 0.858 | 0.225 |
| L CA4 | -0.149 | 0.096 | 0.827 | 0.181 |
| L GC ML DG | -0.142 | 0.096 | 0.827 | 0.197 |
| L Molecular layer | -0.286 | 0.097 | 0.165 | 0.009 |
| L HATA | -0.003 | 0.096 | 0.996 | 0.978 |
| L Fimbria | 0.145 | 0.096 | 0.827 | 0.197 |
| L Hippocampal fissure | -0.067 | 0.096 | 0.975 | 0.557 |
| L Hippocampal tail | 0.097 | 0.096 | 0.975 | 0.400 |
| R Parasubiculum | 0.026 | 0.096 | 0.975 | 0.801 |
| R Presubiculum | 0.054 | 0.096 | 0.975 | 0.635 |
| R Subiculum | -0.044 | 0.096 | 0.975 | 0.652 |
| R CA1 | -0.293 | 0.097 | 0.165 | 0.012 |
| R CA3 | -0.055 | 0.096 | 0.975 | 0.609 |
| R CA4 | -0.032 | 0.096 | 0.975 | 0.763 |
| R Molecular layer | -0.281 | 0.097 | 0.165 | 0.009 |
| R GC ML DG | -0.107 | 0.096 | 0.903 | 0.321 |
| R HATA | -0.039 | 0.096 | 0.975 | 0.652 |
| R Fimbria | 0.210 | 0.097 | 0.443 | 0.053 |
| R Hippocampal fissure | 0.008 | 0.096 | 0.996 | 0.940 |
| R Hippocampal tail | 0.109 | 0.096 | 0.903 | 0.331 |

Abbreviations: cornu ammonis (CA) sectors, CA1, CA2-3, CA4, granule cell layer of dentate gyrus (DG), molecular layer (ML), hippocampus–amygdala transition area (HATA), corticoamygdaloid transition area (CAT), anterior amygdaloid area (AAA).

**Supplemental table S6a:** Mega analysis linear regression results for **bilateral** amygdala and hippocampal subnuclei volumes for OCD patients (n=286) **without MDD comorbidity** compared with adult healthy controls (HCs) (n=338), adjusting for age, sex and scan site and whole volumes for amygdala and hippocampus.

|  | **Cohen’s d**  **(OCD vs HCs)** | **Standard error of *d*** | **FDR corrected**  **p-value** | **Uncorrected p-value** |
| --- | --- | --- | --- | --- |
| Lateral | 0.089 | 0.080 | 0.470 | 0.232 |
| Basal | 0.111 | 0.080 | 0.414 | 0.149 |
| Accessory Basal | 0.023 | 0.080 | 0.854 | 0.757 |
| AAA | -0.055 | 0.080 | 0.580 | 0.414 |
| Central | 0.058 | 0.080 | 0.617 | 0.470 |
| Medial | -0.091 | 0.080 | 0.470 | 0.241 |
| Cortical | -0.014 | 0.080 | 0.892 | 0.853 |
| Corticoamygdaloid transition | -0.009 | 0.080 | 0.892 | 0.892 |
| Paralaminar | 0.135 | 0.080 | 0.320 | 0.076 |
| Parasubiculum | 0.062 | 0.080 | 0.580 | 0.383 |
| Presibiculum | 0.159 | 0.080 | 0.226 | 0.043 |
| Subiculum | 0.060 | 0.080 | 0.580 | 0.387 |
| CA1 | -0.183 | 0.080 | 0.165 | 0.017 |
| CA3 | -0.104 | 0.080 | 0.414 | 0.158 |
| CA4 | -0.117 | 0.080 | 0.382 | 0.109 |
| GC ML DG | -0.167 | 0.080 | 0.165 | 0.024 |
| Molecular layer | -0.176 | 0.080 | 0.165 | 0.020 |
| HATA | -0.064 | 0.080 | 0.491 | 0.281 |
| Fimbria | 0.087 | 0.080 | 0.470 | 0.246 |
| Hippocampal fissure | 0.040 | 0.080 | 0.739 | 0.598 |
| Hippocampal tail | 0.023 | 0.080 | 0.854 | 0.773 |

Abbreviations: cornu ammonis (CA) sectors, CA1, CA2-3, CA4, granule cell layer of dentate gyrus (DG), molecular layer (ML), hippocampus–amygdala transition area (HATA), corticoamygdaloid transition area (CAT), anterior amygdaloid area (AAA).

**Supplemental table S6b:** Mega analysis linear regression results for **left and right** amygdala and hippocampal subnuclei volumes for OCD patients (n=286) **without MDD comorbidity** compared with adult healthy controls (HCs) (n=338), adjusting for age, sex and scan site and whole volumes for amygdala and hippocampus.

|  | **Cohen’s d**  **(OCD vs HCs)** | **Standard error of *d*** | **FDR corrected**  **p-value** | **Uncorrected p-value** |
| --- | --- | --- | --- | --- |
| L Lateral | 0.094 | 0.080 | 0.444 | 0.197 |
| L Basal | 0.106 | 0.080 | 0.444 | 0.170 |
| L Accessory Basal | 0.023 | 0.080 | 0.823 | 0.765 |
| L AAA | -0.035 | 0.080 | 0.803 | 0.622 |
| L Central | 0.131 | 0.080 | 0.368 | 0.105 |
| L Medial | -0.052 | 0.080 | 0.719 | 0.514 |
| L Cortical | 0.035 | 0.080 | 0.803 | 0.647 |
| L Corticoamygdaloid transition | -0.065 | 0.080 | 0.631 | 0.381 |
| L Paralaminar | 0.084 | 0.080 | 0.530 | 0.278 |
| R Lateral | 0.067 | 0.080 | 0.631 | 0.391 |
| R Basal | 0.086 | 0.080 | 0.530 | 0.270 |
| R Accessory Basal | 0.008 | 0.080 | 0.935 | 0.913 |
| R AAA | -0.064 | 0.080 | 0.623 | 0.356 |
| R Central | -0.036 | 0.080 | 0.803 | 0.650 |
| R Medial | -0.113 | 0.080 | 0.444 | 0.149 |
| R Cortical | -0.070 | 0.080 | 0.623 | 0.356 |
| R Corticoamygdaloid transition | 0.043 | 0.080 | 0.719 | 0.510 |
| R Paralaminar | 0.152 | 0.080 | 0.293 | 0.047 |
| L Parasubiculum | 0.000 | 0.080 | 0.996 | 0.996 |
| L Presubiculum | 0.107 | 0.080 | 0.444 | 0.176 |
| L Subiculum | 0.025 | 0.080 | 0.814 | 0.736 |
| L CA1 | -0.143 | 0.080 | 0.293 | 0.056 |
| L CA3 | -0.056 | 0.080 | 0.684 | 0.456 |
| L CA4 | -0.059 | 0.080 | 0.684 | 0.442 |
| L GC ML DG | -0.107 | 0.080 | 0.444 | 0.156 |
| L Molecular layer | -0.151 | 0.080 | 0.293 | 0.049 |
| L HATA | -0.082 | 0.080 | 0.464 | 0.221 |
| L Fimbria | 0.030 | 0.080 | 0.813 | 0.697 |
| L Hippocampal fissure | -0.031 | 0.080 | 0.813 | 0.688 |
| L Hippocampal tail | 0.014 | 0.080 | 0.907 | 0.864 |
| R Parasubiculum | 0.115 | 0.080 | 0.368 | 0.105 |
| R Presubiculum | 0.187 | 0.080 | 0.293 | 0.018 |
| R Subiculum | 0.088 | 0.080 | 0.444 | 0.198 |
| R CA1 | -0.172 | 0.080 | 0.293 | 0.031 |
| R CA3 | -0.133 | 0.080 | 0.342 | 0.073 |
| R CA4 | -0.147 | 0.080 | 0.293 | 0.044 |
| R Molecular layer | -0.157 | 0.080 | 0.293 | 0.034 |
| R GC ML DG | -0.188 | 0.080 | 0.293 | 0.012 |
| R HATA | -0.033 | 0.080 | 0.803 | 0.594 |
| R Fimbria | 0.129 | 0.080 | 0.344 | 0.082 |
| R Hippocampal fissure | 0.097 | 0.080 | 0.444 | 0.201 |
| R Hippocampal tail | 0.026 | 0.080 | 0.814 | 0.736 |

Abbreviations: cornu ammonis (CA) sectors, CA1, CA2-3, CA4, granule cell layer of dentate gyrus (DG), molecular layer (ML), hippocampus–amygdala transition area (HATA), corticoamygdaloid transition area (CAT), anterior amygdaloid area (AAA).

**Supplemental table S7a:** Mega analysis linear regression results for **bilateral** amygdala and hippocampal subnuclei volumes for OCD patients (n=161) **with medication use** compared with adult healthy controls (HCs) (n=291), adjusting for age, sex and scan site and whole volumes for amygdala and hippocampus.

|  | **Cohen’s d**  **(OCD vs HCs)** | **Standard error of *d*** | **FDR corrected**  **p-value** | **Uncorrected p-value** |
| --- | --- | --- | --- | --- |
| **Lateral** | **0.233** | **0.095** | **0.049** | **0.008** |
| **Basal** | **0.250** | **0.095** | **0.049** | **0.009** |
| Accessory Basal | -0.010 | 0.094 | 0.914 | 0.914 |
| AAA | 0.084 | 0.094 | 0.427 | 0.289 |
| Central | 0.171 | 0.094 | 0.187 | 0.084 |
| Medial | -0.109 | 0.094 | 0.427 | 0.251 |
| Cortical | -0.072 | 0.094 | 0.495 | 0.448 |
| Corticoamygdaloid transition | -0.088 | 0.094 | 0.427 | 0.310 |
| Paralaminar | 0.226 | 0.095 | 0.069 | 0.019 |
| Parasubiculum | -0.079 | 0.094 | 0.427 | 0.355 |
| Presibiculum | 0.089 | 0.094 | 0.427 | 0.351 |
| Subiculum | 0.079 | 0.094 | 0.427 | 0.366 |
| CA1 | -0.093 | 0.094 | 0.427 | 0.323 |
| CA3 | -0.153 | 0.094 | 0.187 | 0.089 |
| CA4 | -0.194 | 0.095 | 0.085 | 0.032 |
| **GC ML DG** | **-0.264** | **0.095** | **0.042** | **0.004** |
| **Molecular layer** | **-0.296** | **0.095** | **0.042** | **0.002** |
| HATA | -0.164 | 0.094 | 0.069 | 0.020 |
| Fimbria | 0.191 | 0.094 | 0.085 | 0.031 |
| Hippocampal fissure | 0.126 | 0.094 | 0.314 | 0.165 |
| Hippocampal tail | -0.051 | 0.094 | 0.626 | 0.596 |

Abbreviations: cornu ammonis (CA) sectors, CA1, CA2-3, CA4, granule cell layer of dentate gyrus (DG), molecular layer (ML), hippocampus–amygdala transition area (HATA), corticoamygdaloid transition area (CAT), anterior amygdaloid area (AAA).

**Supplemental table S7b:** Mega analysis linear regression results for **left and right** amygdala and hippocampal subnuclei volumes for OCD patients (n=161) **with medication use** compared with adult healthy controls (HCs) (n=291), adjusting for age, sex and scan site and whole volumes for amygdala and hippocampus.

|  | **Cohen’s d**  **(OCD vs HCs)** | **Standard error of *d*** | **FDR corrected**  **p-value** | **Uncorrected p-value** |
| --- | --- | --- | --- | --- |
| L Lateral | 0.162 | 0.094 | 0.154 | 0.056 |
| L Basal | 0.212 | 0.095 | 0.108 | 0.027 |
| L Accessory Basal | -0.006 | 0.094 | 0.974 | 0.951 |
| L AAA | 0.031 | 0.094 | 0.803 | 0.708 |
| L Central | 0.237 | 0.095 | 0.103 | 0.017 |
| L Medial | -0.137 | 0.094 | 0.318 | 0.159 |
| L Cortical | -0.050 | 0.094 | 0.743 | 0.602 |
| L Corticoamygdaloid transition | -0.143 | 0.094 | 0.286 | 0.129 |
| L Paralaminar | 0.182 | 0.094 | 0.154 | 0.060 |
| R Lateral | 0.262 | 0.095 | 0.077 | 0.005 |
| R Basal | 0.220 | 0.095 | 0.108 | 0.024 |
| R Accessory Basal | -0.029 | 0.094 | 0.844 | 0.764 |
| R AAA | 0.121 | 0.094 | 0.300 | 0.143 |
| R Central | 0.058 | 0.094 | 0.701 | 0.551 |
| R Medial | -0.065 | 0.094 | 0.678 | 0.500 |
| R Cortical | -0.097 | 0.094 | 0.499 | 0.318 |
| R Corticoamygdaloid transition | -0.018 | 0.094 | 0.886 | 0.823 |
| R Paralaminar | 0.211 | 0.095 | 0.108 | 0.028 |
| L Parasubiculum | -0.088 | 0.094 | 0.499 | 0.336 |
| L Presubiculum | 0.124 | 0.094 | 0.361 | 0.198 |
| L Subiculum | 0.092 | 0.094 | 0.499 | 0.323 |
| L CA1 | -0.102 | 0.094 | 0.461 | 0.264 |
| L CA3 | -0.124 | 0.094 | 0.346 | 0.181 |
| L CA4 | -0.175 | 0.094 | 0.154 | 0.062 |
| L GC ML DG | -0.224 | 0.095 | 0.103 | 0.017 |
| L Molecular layer | -0.260 | 0.095 | 0.077 | 0.006 |
| L HATA | -0.157 | 0.094 | 0.154 | 0.052 |
| L Fimbria | 0.090 | 0.094 | 0.499 | 0.338 |
| L Hippocampal fissure | 0.010 | 0.094 | 0.966 | 0.920 |
| L Hippocampal tail | 0.002 | 0.094 | 0.987 | 0.987 |
| R Parasubiculum | -0.042 | 0.094 | 0.743 | 0.624 |
| R Presubiculum | 0.045 | 0.094 | 0.743 | 0.637 |
| R Subiculum | 0.057 | 0.094 | 0.678 | 0.498 |
| R CA1 | -0.062 | 0.094 | 0.694 | 0.529 |
| R CA3 | -0.159 | 0.094 | 0.180 | 0.077 |
| R CA4 | -0.178 | 0.094 | 0.154 | 0.050 |
| R Molecular layer | -0.259 | 0.095 | 0.077 | 0.009 |
| R GC ML DG | -0.249 | 0.095 | 0.077 | 0.008 |
| R HATA | -0.140 | 0.094 | 0.154 | 0.051 |
| R Fimbria | 0.253 | 0.095 | 0.077 | 0.004 |
| R Hippocampal fissure | 0.206 | 0.095 | 0.108 | 0.024 |
| R Hippocampal tail | -0.088 | 0.094 | 0.499 | 0.345 |

Abbreviations: cornu ammonis (CA) sectors, CA1, CA2-3, CA4, granule cell layer of dentate gyrus (DG), molecular layer (ML), hippocampus–amygdala transition area (HATA), corticoamygdaloid transition area (CAT), anterior amygdaloid area (AAA).

**Supplemental table S8a:** Mega analysis linear regression results for **bilateral** amygdala and hippocampal subnuclei volumes for OCD patients (n=220) **without medication use** compared with adult healthy controls (HCs) (n=338), adjusting for age, sex and scan site and whole volumes for amygdala and hippocampus.

|  | **Cohen’s d**  **(OCD vs HCs)** | **Standard error of *d*** | **FDR corrected**  **p-value** | **Uncorrected p-value** |
| --- | --- | --- | --- | --- |
| Lateral | -0.016 | 0.085 | 0.899 | 0.841 |
| Basal | -0.028 | 0.085 | 0.899 | 0.732 |
| Accessory Basal | -0.017 | 0.085 | 0.899 | 0.834 |
| AAA | -0.141 | 0.085 | 0.252 | 0.048 |
| Central | -0.088 | 0.085 | 0.816 | 0.311 |
| Medial | -0.070 | 0.085 | 0.856 | 0.408 |
| Cortical | 0.012 | 0.085 | 0.899 | 0.882 |
| Corticoamygdaloid transition | 0.017 | 0.085 | 0.899 | 0.817 |
| Paralaminar | 0.073 | 0.085 | 0.856 | 0.371 |
| Parasubiculum | 0.142 | 0.085 | 0.258 | 0.061 |
| Presibiculum | 0.195 | 0.085 | 0.235 | 0.022 |
| Subiculum | 0.017 | 0.085 | 0.899 | 0.820 |
| CA1 | -0.281 | 0.085 | 0.017 | 0.001 |
| CA3 | -0.038 | 0.085 | 0.899 | 0.636 |
| CA4 | -0.045 | 0.085 | 0.899 | 0.567 |
| GC ML DG | -0.085 | 0.085 | 0.816 | 0.280 |
| Molecular layer | -0.164 | 0.085 | 0.252 | 0.039 |
| HATA | 0.008 | 0.085 | 0.899 | 0.899 |
| Fimbria | 0.041 | 0.085 | 0.899 | 0.612 |
| Hippocampal fissure | -0.037 | 0.085 | 0.899 | 0.657 |
| Hippocampal tail | 0.147 | 0.085 | 0.292 | 0.083 |

Abbreviations: cornu ammonis (CA) sectors, CA1, CA2-3, CA4, granule cell layer of dentate gyrus (DG), molecular layer (ML), hippocampus–amygdala transition area (HATA), corticoamygdaloid transition area (CAT), anterior amygdaloid area (AAA).

**Supplemental table S8b:** Mega analysis linear regression results for **left and right amygdala** and hippocampal subnuclei volumes for OCD patients (n=220) **without medication use** compared with adult healthy controls (HCs) (n=338), adjusting for age, sex and scan site and whole volumes for amygdala and hippocampus.

|  | **Cohen’s d**  **(OCD vs HCs)** | **Standard error of *d*** | **FDR corrected**  **p-value** | **Uncorrected p-value** |
| --- | --- | --- | --- | --- |
| L Lateral | -0.009 | 0.085 | 0.989 | 0.908 |
| L Basal | -0.007 | 0.085 | 0.989 | 0.935 |
| L Accessory Basal | -0.002 | 0.085 | 0.989 | 0.983 |
| L AAA | -0.122 | 0.085 | 0.430 | 0.102 |
| L Central | -0.019 | 0.085 | 0.989 | 0.831 |
| L Medial | -0.013 | 0.085 | 0.989 | 0.880 |
| L Cortical | 0.053 | 0.085 | 0.989 | 0.521 |
| L Corticoamygdaloid transition | -0.007 | 0.085 | 0.989 | 0.929 |
| L Paralaminar | 0.034 | 0.085 | 0.989 | 0.680 |
| R Lateral | -0.012 | 0.085 | 0.989 | 0.885 |
| R Basal | -0.041 | 0.085 | 0.989 | 0.619 |
| R Accessory Basal | -0.028 | 0.085 | 0.989 | 0.733 |
| R AAA | -0.129 | 0.085 | 0.411 | 0.082 |
| R Central | -0.147 | 0.085 | 0.411 | 0.088 |
| R Medial | -0.106 | 0.085 | 0.692 | 0.214 |
| R Cortical | -0.027 | 0.085 | 0.989 | 0.742 |
| R Corticoamygdaloid transition | 0.038 | 0.085 | 0.989 | 0.582 |
| R Paralaminar | 0.094 | 0.085 | 0.743 | 0.248 |
| L Parasubiculum | 0.066 | 0.085 | 0.977 | 0.419 |
| L Presubiculum | 0.150 | 0.085 | 0.411 | 0.081 |
| L Subiculum | -0.022 | 0.085 | 0.989 | 0.776 |
| L CA1 | -0.188 | 0.085 | 0.218 | 0.021 |
| L CA3 | 0.001 | 0.085 | 0.989 | 0.989 |
| L CA4 | -0.003 | 0.085 | 0.989 | 0.968 |
| L GC ML DG | -0.040 | 0.085 | 0.989 | 0.618 |
| L Molecular layer | -0.122 | 0.085 | 0.522 | 0.137 |
| L HATA | -0.017 | 0.085 | 0.989 | 0.812 |
| L Fimbria | 0.022 | 0.085 | 0.989 | 0.795 |
| L Hippocampal fissure | -0.046 | 0.085 | 0.989 | 0.584 |
| L Hippocampal tail | 0.086 | 0.085 | 0.894 | 0.321 |
| R Parasubiculum | 0.177 | 0.085 | 0.218 | 0.020 |
| R Presubiculum | 0.205 | 0.085 | 0.218 | 0.017 |
| R Subiculum | 0.052 | 0.085 | 0.989 | 0.475 |
| **R CA1** | **-0.298** | **0.085** | **0.024** | **0.001** |
| R CA3 | -0.070 | 0.085 | 0.945 | 0.382 |
| R CA4 | -0.074 | 0.085 | 0.894 | 0.340 |
| R Molecular layer | -0.166 | 0.085 | 0.236 | 0.034 |
| R GC ML DG | -0.112 | 0.085 | 0.573 | 0.164 |
| R HATA | 0.030 | 0.085 | 0.989 | 0.643 |
| R Fimbria | 0.059 | 0.085 | 0.989 | 0.457 |
| R Hippocampal fissure | -0.017 | 0.085 | 0.989 | 0.835 |
| R Hippocampal tail | 0.177 | 0.085 | 0.236 | 0.033 |

Abbreviations: cornu ammonis (CA) sectors, CA1, CA2-3, CA4, granule cell layer of dentate gyrus (DG), molecular layer (ML), hippocampus–amygdala transition area (HATA), corticoamygdaloid transition area (CAT), anterior amygdaloid area (AAA).

**Supplemental table 9:** Mega analysis linear regression results for the association between amygdala and hippocampal subnuclei volumes in OCD patients (n=351) and **YBOCS score**.

|  | **rME** | **Standard error** | **FDR corrected**  **p-value** | **Uncorrected p-value** |
| --- | --- | --- | --- | --- |
| Lateral | 0.023 | 0.054 | 0.760 | 0.615 |
| Basal | 0.058 | 0.054 | 0.711 | 0.221 |
| Accessory Basal | 0.074 | 0.054 | 0.711 | 0.112 |
| AAA | 0.018 | 0.054 | 0.773 | 0.662 |
| Central | 0.048 | 0.054 | 0.757 | 0.360 |
| Medial | 0.055 | 0.054 | 0.711 | 0.254 |
| Cortical | 0.058 | 0.054 | 0.711 | 0.218 |
| Corticoamygdaloid transition | -0.015 | 0.054 | 0.802 | 0.726 |
| Paralaminar | 0.002 | 0.054 | 0.964 | 0.964 |
| Parasubiculum | -0.048 | 0.054 | 0.711 | 0.305 |
| Presibiculum | -0.057 | 0.054 | 0.711 | 0.282 |
| Subiculum | -0.024 | 0.054 | 0.760 | 0.606 |
| CA1 | 0.051 | 0.054 | 0.711 | 0.305 |
| CA3 | 0.028 | 0.054 | 0.760 | 0.565 |
| CA4 | 0.035 | 0.054 | 0.760 | 0.473 |
| GC ML DG | 0.037 | 0.054 | 0.760 | 0.440 |
| Molecular layer | 0.025 | 0.054 | 0.760 | 0.615 |
| HATA | 0.002 | 0.054 | 0.964 | 0.959 |
| Fimbria | -0.034 | 0.054 | 0.760 | 0.492 |
| Hippocampal fissure | -0.085 | 0.054 | 0.711 | 0.089 |
| Hippocampal tail | -0.083 | 0.054 | 0.711 | 0.113 |

Abbreviations: cornu ammonis (CA) sectors, CA1, CA2-3, CA4, granule cell layer of dentate gyrus (DG), molecular layer (ML), hippocampus–amygdala transition area (HATA), corticoamygdaloid transition area (CAT), anterior amygdaloid area (AAA).
